# Inverted Alu repeats in loop-out exon skipping across hominoid evolution

**DOI:** 10.1101/2025.03.07.642063

**Authors:** Danielle Denisko, Jeonghyeon Kim, Jayoung Ku, Boxun Zhao, Eunjung Alice Lee

**Affiliations:** Division of Genetics and Genomics, Boston Children’s Hospital and Harvard Medical School, Boston, MA 02115, USA; Department of Biomedical Informatics, Harvard Medical School, Boston, MA 02115, USA; Department of Chemical and Systems Biology, Stanford University, Stanford, CA 94305, USA; Broad Institute of MIT and Harvard, Cambridge, MA 02115, USA; Manton Center for Orphan Disease Research, Boston Children’s Hospital, Boston, MA 02115, USA

**Author notes:** Corresponding author: (B.Z.) and (E.A.L.). Equal contribution (Danielle Denisko, Jeonghyeon Kim).

**Keywords:** Exon skipping, Alternative splicing, Transposable elements, Alu, Inverted repeats, Hominoid evolution

## Abstract

**Background:** Changes in RNA splicing over the course of evolution have profoundly diversified the functional landscape of the human genome. While DNA sequences proximal to intron-exon junctions are known to be critical for RNA splicing, the impact of distal intronic sequences remains underexplored. Emerging evidence suggests that inverted pairs of intronic Alu elements can promote exon skipping by forming RNA stem-loop structures. However, their prevalence and influence throughout evolution remain unknown.

**Results:** Here, we present a systematic analysis of inverted Alu pairs across the human genome to assess their impact on exon skipping through predicted RNA stem-loop formation and their relevance to hominoid evolution. We found that inverted Alu pairs, particularly pairs of AluY-AluSx1 and AluSz-AluSx, are enriched in the flanking regions of skippable exons genome-wide and are predicted to form stable stem-loop structures. Exons defined by weak 3′ acceptor and strong 5′ donor splice sites appear especially prone to this skipping mechanism. Through comparative genome analysis across nine primate species, we identified 67,126 hominoid-specific Alu insertions, primarily from AluY and AluS subfamilies, which form inverted pairs enriched across skippable exons in genes of ubiquitination-related pathways. Experimental validation of exon skipping among several hominoid-specific inverted Alu pairs further reinforced their potential evolutionary significance.

**Conclusion:** This work extends our current knowledge of the roles of RNA secondary structure formed by inverted Alu pairs and details a newly emerging mechanism through which transposable elements have contributed to genomic innovation across hominoid evolution at the transcriptomic level.

## Introduction

Acting on the vast majority of mammalian multi-exon genes, alternative splicing has increased transcriptomic and proteomic diversity across species throughout evolution [1–4]. Splice site determination is orchestrated by the assembly of the spliceosome, mediated by core splicing signals, as well as splicing enhancer and silencer motifs and their cognate binding factors [4,5]. Genomic and epigenomic changes influence alternative splicing decisions by disrupting or creating splice sites and modulating splice factor binding, together revealing the complex combinatorial nature underlying alternative splicing regulatory networks [6–9]. The most common form of alternative splicing in metazoans is exon skipping, which can be described as the exclusion of an exon from a mature RNA transcript [10]. In primates, alternative splicing networks have evolved to increase differential exon usage, positioning isoform regulation as a crucial evolutionary mechanism that expands the cellular regulatory and functional repertoire, likely contributing to phenotypic complexity [11–13].

While splicing outcomes associated with sequences proximal to intron-exon junctions have been extensively studied, less is known about the importance of more distal sequences. RNA secondary structure offers an avenue through which distal sequences may impart their effects. Considering a large portion of splicing occurs co-transcriptionally, the secondary structure of nascent RNA has the ability to influence splice site usage by either sequestering motifs within base-paired double-stranded stem regions, hindering access to splicing machinery, or conversely by facilitating the recognition of these motifs by presenting them in more accessible, single-stranded loop regions of the transcript [14–16]. While investigations into the impact of stem-loop structures on splicing have generally focused on structures modulating individual splice regulatory site accessibility, evidence suggests that RNA secondary structure also mediates an unconventional mechanism of exon skipping [17]. Coined as “jump splicing” in the 1980s, loops encompassing exons within RNA transcripts were found to be able to promote exon skipping under certain conditions [18,19]. More recent studies have also correlated circular RNA (circRNA) biogenesis with exon skipping, further supporting the existence of a related loop-out mechanism [20].

An overarching theme of the proposed loop-out exon skipping mechanism is the presence of reverse complementary sequences flanking each side of such skippable exons. Making up about half of the human genome, repetitive sequence elements called transposons account for a large fraction of reverse complementary sequences found genome-wide [21]. Retrotransposons form a distinct class of transposons that insert copies of themselves into the genome through an RNA intermediate in a “copy and paste” manner [22]. Among retrotransposons, only long interspersed element-1 (L1), Alu, and SINE-R/VNTR/Alu (SVA) elements have remained active throughout primate evolution up to present-day humans [23], together orchestrating an extensive fossil record of genomic innovation.

Alu elements constitute a particular type of short (approximately 300 bp), primate-specific retrotransposons that are prone to forming double-stranded RNA stem-loop structures through base pairing interactions with neighboring Alus lying in inverted orientation [24,25]. There are three main Alu subfamilies, differing based on the mutations amassed in their underlying consensus sequences: AluJ elements form the oldest dimeric subfamily of Alus, having appeared approximately 80 million years ago, while AluS elements are considered to be 30-50 million years old, and AluY elements form the youngest subfamily at less than 15 million years old [26–28]. Alus have accumulated in primate genomes over time largely through genetic drift [29–32], while negative selection has acted to remove their deleterious effects. Beyond shaping primate genomes through insertional mutagenesis, Alu elements have also driven genome instability by serving as substrates for non-allelic homologous recombination events and have increased isoform diversity through exonization [26,33,34]. Currently, the human genome contains over 1.2 million fixed copies of Alus, with up to 75% of genes containing at least one copy [35,36]. Alu insertions are also present as polymorphisms across the human population [37,38], and have been implicated as drivers of several genetic diseases [39–42].

Due to their biased localization toward gene-rich regions, many Alus are clustered closely together, with nearly half located in introns [24,43]. Their proximity and high sequence similarity enable them to form inverted Alu pair RNA stem-loop structures, which play crucial roles in a wide range of biological processes spanning from gene regulation to immunology [44]. Inverted Alu stem-loops have also been causally linked to disease, such as severe infantile isolated exocrine pancreatic insufficiency [45]. Recently, a new role for inverted Alus consisting of the mediation of loop-out exon skipping has been discovered. Specifically, evidence suggests that a hominoid-specific AluY insertion forms an inverted pair with a pre-existing Alu element in tail development gene *TBXT*, arbitrating an exon skipping isoform that may underlie the loss of tails in hominoid species [46]. This exciting discovery raises important questions about the prevalence of an inverted Alu-mediated exon skipping mechanism and its impact on primate evolution.

In this work, we undertake the first systematic assessment of the contribution of fixed Alu repeat pairs in mediating loop-out exon skipping across the human genome. Through comparative genomic analysis, we further characterize the contributions of hominoid-specific inverted Alu repeats in exon skipping to better understand the impact of repeat pairs on hominoid evolution.

## Results

### Inverted repeat Alu pairs are enriched in the flanks of skippable exons genome-wide

Given the potential abundance of inverted Alus lying across spliced exons that may form base pair interactions, we first sought to characterize properties of such Alu pairings genome-wide using a systematic and unbiased approach. For this, we identified all possible pairs of Alu elements spanning skippable and constitutive exons across the human reference genome using a maximum window size of 5,000 bp (Figure 1A). We defined skippable and constitutive exon sets according to HEXEvent [47] and ExonSkipDB [48] databases. In total, we extracted 26,803 skippable and 23,540 constitutive exons, resulting in 347,816 and 232,493 Alu pairs respectively flanking each exon type. Across all flanking window sizes ranging from 500 bp to 5,000 bp, we found that inverted Alu pairs were significantly enriched in the flanks of skippable exons compared to constitutive exons (adjusted *p* < 0.001, Fisher’s exact test; Figure 1B and Figure S1A). Inverted Alu pairs, however, were not significantly enriched within adjacent, non-overlapping windows, indicating that symmetry of the paired Alu elements relative to the exon boundaries is unlikely to be crucial for skipping (Figure S1B).

**Figure 1.**
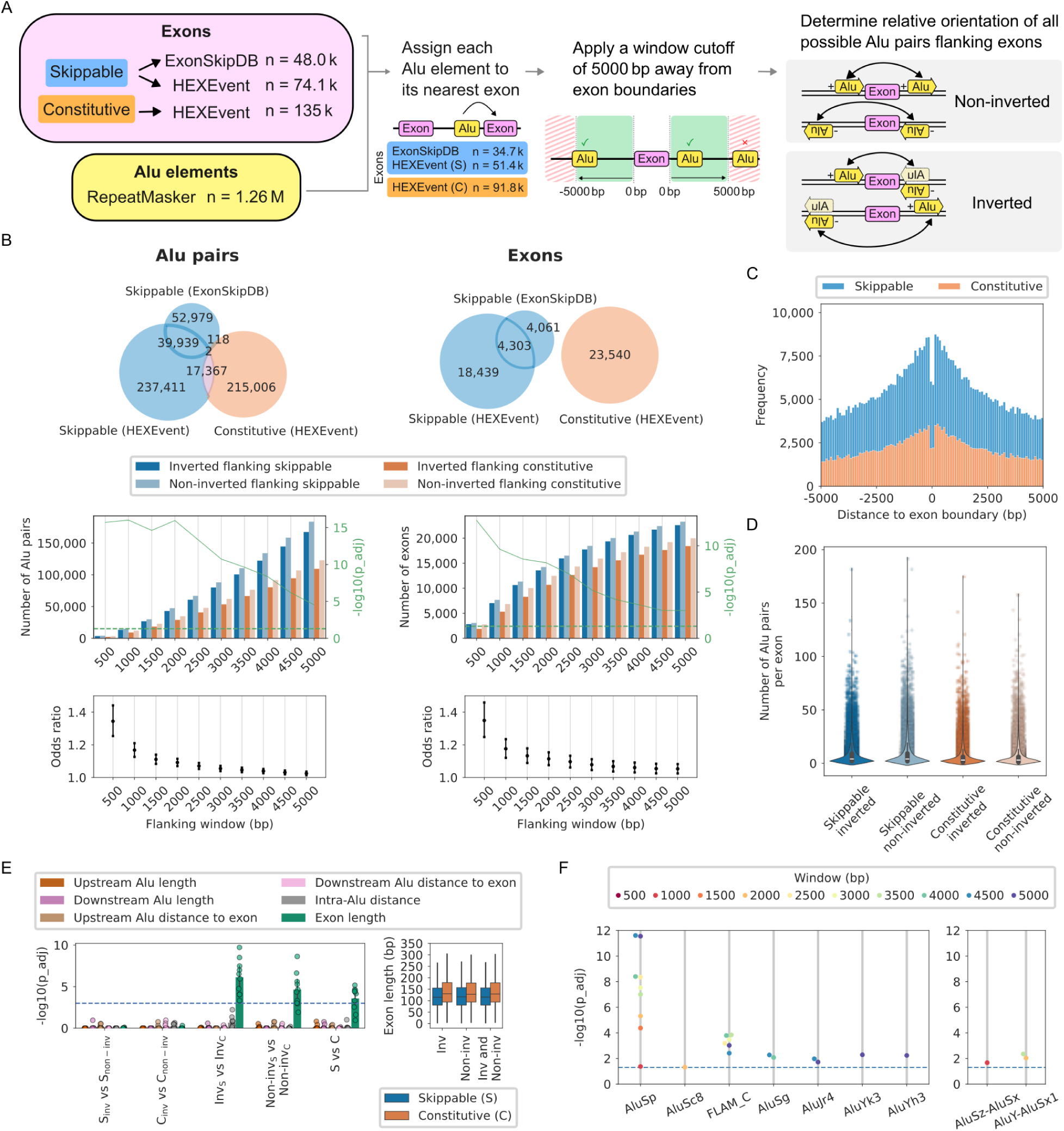
Inverted repeat Alu pairs are enriched in the flanks of skippable exons genome-wide. **(A)** Method to identify all possible fixed Alu pairs flanking skippable and constitutive exons in the human genome within windows of up to +/− 5000 bp surrounding exons. S: skippable, C: constitutive. **(B)** Top: Number of Alu pairs (left) and exons (right) per exon database: HEXEvent and ExonSkipDB. Middle: Enrichment of inverted vs. non-inverted Alu pairs flanking skippable exons, compared to those flanking constitutive exons. Fisher’s exact (FE) test on the number of Alu pairs (left) and exons flanked by Alu pairs (right) across windows of size 500 bp to 5000 bp. Bonferroni-corrected one-tailed FE test p-values. Bottom: Odds ratios with 95% confidence intervals corresponding to the middle panel. **(C)** The distance of each Alu in a pair to its flanked exon (bp). **(D)** The number of flanking Alu pairs per exon within 5000 bp flanking windows. **(E)** A comparison of distance and length features across exon types and Alu pair inversion. 10-fold random subsampling of 1000 pairs of Alus within 1000 bp flanking windows. Distances and lengths are measured in bp. Bonferroni-corrected Kolmogorov–Smirnov test p-values. S: skippable, C: constitutive, Inv: inverted, Non-inv: non-inverted. **(F)** Individual Alu (left) and pairs of Alu (right) subfamilies enriched in inverted orientation flanking skippable exons. Bonferroni-corrected one-tailed FE test p-values.

Fixed Alus were generally 300 bp in length, with a smaller peak of truncated elements at around 150-160 bp, in accordance with previous reports (Figures S1C and S1D) [49,50]. The distance of Alu elements within pairs relative to exon boundaries recapitulated the known trend of negative purifying selection occurring within ∼100 bp of the exon boundaries (Figure 1C and Figure S1E) [51,52]. To assess whether individual exons may bias the enrichment tests, we confirmed that the number of Alu pairs per exon was comparable across all exon types and pair inversion combinations (Figure 1D).

We next sought to understand whether individual length and distance features of the underlying elements differed across exon types and Alu pair inversion. Using a 10-fold downsampling scheme to adjust for high statistical power, we compared the upstream and downstream Alu lengths, the distance of the upstream and downstream Alus to the intervening exon boundaries, the distance between Alus, and the length of the exon for each sampled Alu pair (see Methods). Across all comparisons, we found that only the exon length differed significantly between skippable and constitutive exon groups, which we observed for both inverted and non-inverted pair configurations and across all tested flanking windows of size 1,000 bp, 2,000 bp, and 5,000 bp (average *p* < 0.05, Bonferroni-adjusted Kolmogorov–Smirnov test; Figure 1E and Figure S1F). More specifically, we observed that skippable exons tend to be shorter than constitutive exons, a result that aligns with previous findings for alternative exons in general [53,54].

Since individual Alu subfamilies arose at different points throughout primate evolution, their underlying sequences have diverged according to accumulated mutations. Considering that the degree of homology determines the strength of base pairing between adjacent repetitive sequences in the genome, we next examined the enrichment of Alu subfamilies within pairs. Overall, we found that several subfamilies across AluJ, AluS, and AluY were significantly enriched as individual members within inverted pairs flanking skippable exons (adjusted *p* < 0.05, Fisher’s exact test; Figure 1F). Most notably, AluSp elements were significantly enriched across nearly all flanking windows spanning from 500 bp to 5,000 bp. In terms of specific Alu subfamily pairings, we found two pairings that were significantly enriched: AluSz-AluSx in 1,000 bp windows (adjusted *p* = 0.021), and AluY-AluSx1 in both 2,000 bp (adjusted *p* = 0.0092) and 3,500 bp (adjusted *p* = 0.0044) windows. Intriguingly, it was a pairing between a young AluY element and an ancestral AluSx1 element that was identified in the aforementioned study as a genetic driver of hominoid tail-loss through the proposed loop-out exon skipping mechanism [46]. The underlying Alu element count for each subfamily was comparable across groups, suggesting that these findings did not arise from inflated counts (Figure S1G). Taken together, our initial overview of the human Alu pair landscape reveals that inverted Alu pairs across Alu lineages are enriched in the flanks of skippable exons genome-wide suggesting their potential widespread involvement in mediating exon skipping.

### Inverted Alu pairs flanking skipped exons are predicted to form more stable RNA secondary structures, especially among exons with weak exon definition

To understand how structure may play a role in exon skipping, we next studied the predicted RNA secondary structure of Alu pairs. For this, we selected Alu pairs falling within flanking windows of size 1,000 bp, which demonstrated significant enrichment of inverted Alu pairs across skippable exons (Figure 1B). For pairs falling within this window, we extracted sequences spanning from the start of the upstream Alu to the end of the downstream Alu, through the intervening flanked exon (Figure 2A), using random size-matched genic regions as controls. We predicted RNA secondary structures and their associated minimum free energies (MFEs) for 49,891 Alu pair sequences using RNAfold [55,56]. Of these, most pairs flanked skippable exons (n = 28,620) compared to constitutive (n = 21,271) exons, while there were fewer inverted (n = 22,619) compared to non-inverted (n = 27,272) pairs. To account for the known bias of longer sequences forming more stable structures, we adjusted MFE scores by length (Figure 2B). Overall, we found that inverted Alu pairs were predicted to form much more stable structures as compared to non-inverted Alu pairs (*d* = −1.25) as well as random genic region controls (*d* = −2.18 and *d* = −2.08), suggesting their ability to form stem-loop structures (Figure 2C). Non-inverted pairs were also predicted to form structures with much lower MFE than random genic sequences (*d* = −0.98 and *d* = −0.88), although the magnitude of the effect was smaller than observed for inverted pairs. Surprisingly, we observed only a small difference in MFE between skippable and constitutive exon sets among inverted Alu pairs (*d* = −0.12), which was comparable to non-inverted pairs (*d* = −0.13). While we did not expect to observe a large difference in MFE across the overall sets of Alu pair sequences flanking skippable and constitutive exons, observing a small effect size specifically among inverted Alu pairs suggests that loop-out exon skipping is not the only major mechanism mediating exon skipping in the presence of inverted Alu pairs. Rather, other genomic or epigenomic determinants may take precedence over RNA secondary structure folding dictated by inverted Alu pairs and govern alternative splicing outcomes.

**Figure 2.**
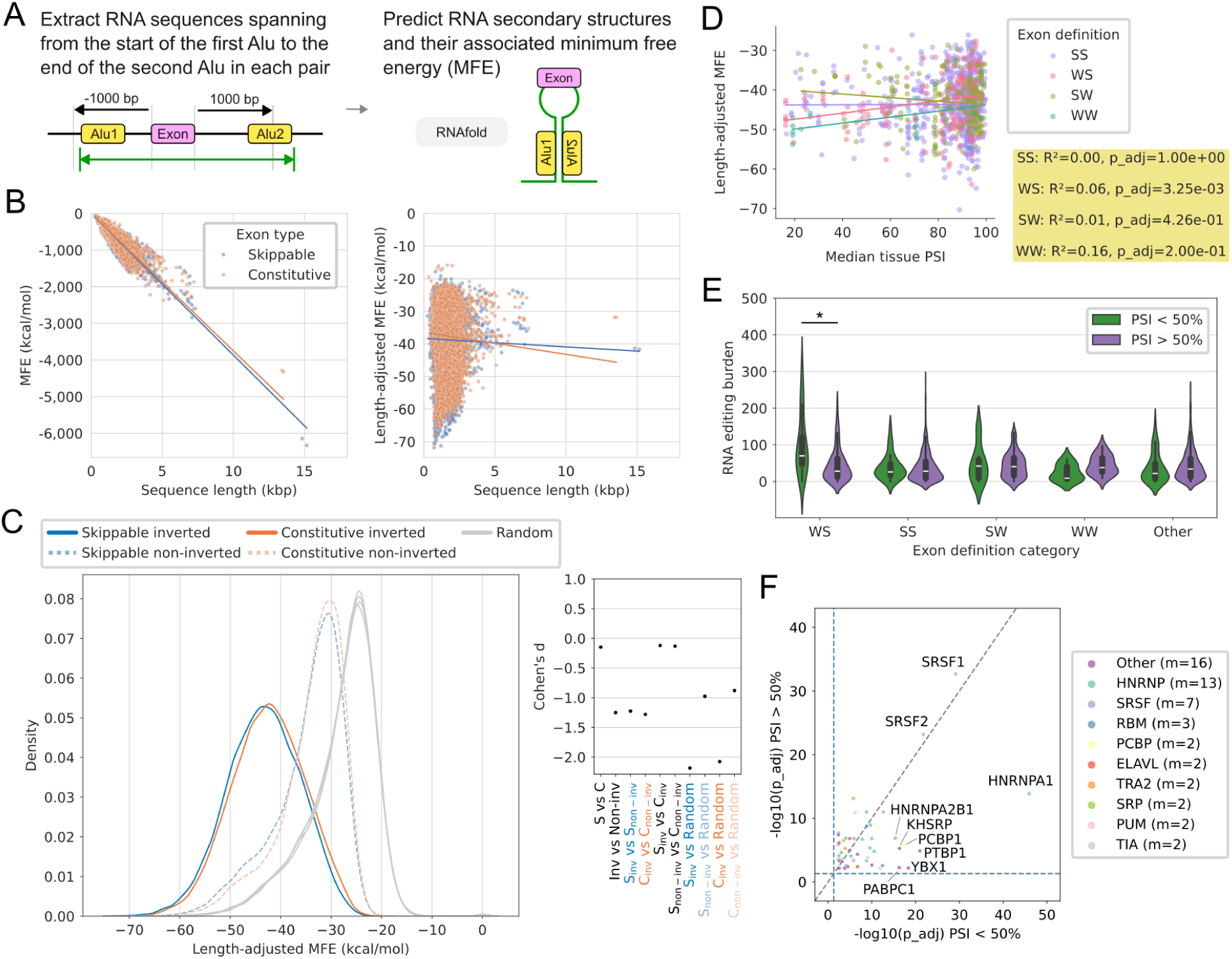
Inverted repeat Alus flanking skippable exons show a greater propensity to form stable RNA secondary structures. **(A)** Method to extract sequences for RNA secondary structure prediction. Strand-specific sequences spanning from the start of the upstream Alu to the end of the downstream Alu of pairs lying within +/−1000 bp windows are passed into RNAfold. **(B)** Raw minimum free energy (MFE) scores (left) are adjusted according to their lengths (right) to remove length bias. **(C)** MFE distributions across exon type and Alu inversion categories and their associated effect sizes as measured by Cohen’s d. **(D)** Correlation between the median percent spliced-in (PSI) reported across 56 tissues in the ASCOT database and the adjusted MFE across exons of different strengths, among skippable exons flanked by inverted Alu pairs. SS: strong 3’ acceptor, strong 5’ donor; WS: weak 3’ acceptor, strong 5’ donor; SW: strong 3’ acceptor, weak 5’ donor; WW: weak 3’ acceptor, weak 5’ donor. Bonferroni-adjusted linear regression p-values. **(E)** RNA editing burden from the REDIportal across exon definition categories among skippable exons flanked by inverted Alu pairs. Two-sided Student’s t-test Bonferroni-adjusted p-values. *p < 0.001. **(F)** Motif enrichment analysis of inverted Alu pairs flanking skippable exons defined by a weak 3’ acceptor and a strong 5’ donor splice site (WS). Differential motif analysis was performed using SEA with either PSI < 50% (x-axis) or PSI > 50% (y-axis) sequences as primary input sequences, and the other set as control sequences. P-values were calculated by SEA using Fisher’s exact test and combined across multiple motif instances using Fisher’s method. P-values were adjusted using the Benjamini–Hochberg procedure. Blue dashed lines indicate p_adj = 0.05.

To further investigate the loop-out mechanism in exon skipping, we next hypothesized that loop-out exon skipping is employed by exons marked by a weak exon definition. It has previously been established that exons marked by weak exon definition are generally more likely to undergo alternative splicing than those marked by strong exon definition [54]. Here, we similarly reasoned that in the absence of strong 3′ acceptor or 5′ donor splice sites, typical splice regulatory factors recognizing these sites would be less likely to bind, thereby allowing other factors involved in the promotion of looping to bind. To test this hypothesis, we split exons into groups according to their exon definition, with weak (W) splice sites having a MaxEntScan score less than 6, and strong (S) splice sites having a score greater than 8 [57,58]. We assigned combined labels to each exon corresponding to the acceptor followed by the donor splice site strengths (for example, “WS” corresponds to an exon with a weak 3′ acceptor site and a strong 5′ donor site). We further annotated each exon with the degree of exon skipping as measured by percent spliced-in (PSI) values across 56 tissues from the ASCOT catalog of alternative splicing events [59]. This left us with 3,405 Alu pairs flanking skippable exons, among which 1,615 of these pairs were inverted. Among inverted pairs flanking skippable exons, most sequences contained exons designated as SS (n = 639), followed by SW (n = 184), WS (n = 177), and WW (n = 24), while the rest (n = 591) did not fall within those categories. Under the expectation that stem-loop structures promote exon skipping, we postulated that exons making use of loop-out skipping should demonstrate a positive correlation between their degree of exclusion and their structural stability. Plotting MFE against PSI, we found that among inverted Alu pairs flanking skippable exons, only those flanking WS exons demonstrated a statistically significant positive correlation (slope = 0.08, R² = 0.06, adjusted *p* = 0.0033; Figure 2D and Figure S2B). Although this correlation is weak, this could be explained by the involvement of other layers of splicing regulation.

Since RNA stem-loops form the substrate for ADAR enzymes, we next assessed the underlying RNA editing burden [60]. Comparing RNA editing burden across exon definition groups of inverted Alus flanking skippable exons, we found that only WS exons demonstrated a significantly higher RNA editing burden among exons that were more frequently skipped (PSI < 50%) compared to exons that were more frequently retained (PSI > 50%; adjusted *p* = 2.9e-6; Figure 2E). These findings corroborate the relevance of loop-out exon skipping among exons with a weak 3′ acceptor site and a strong 5′ donor site.

To elucidate potential factors involved in mediating loop-out exon skipping, we next performed motif enrichment analysis on inverted Alu pairs flanking skippable WS exons that have a higher tendency to be skipped (median tissue PSI < 50%; n = 25) relative to those that tend to be retained (median tissue PSI > 50%; n = 152). We observed an enrichment of several-splicing related factors in the Alu pair sequence set of those flanking exons with a higher tendency for skipping, including PTBP1, PCBP1, hnRNPA2B1, and hnRNPA1 (Figure 2F) [61–64]. Additionally, some factors such as PTBP1, hnRNPA2B1, hnRNPA1, and KHSRP are implicated in circRNA metabolism [64–67]. Interestingly, hnRNPA1, which showed the most striking enrichment in sequences with higher exon skipping tendency, has been previously linked to exon skipping through a loop-out mechanism [64]. Collectively, our results suggest that inverted Alus may play a role in mediating exon skipping in the absence of a strong 3′ acceptor splice site, facilitated by factors previously implicated in looping mechanisms as well as factors that have not yet been explored in this context.

### Skipped exons flanked by inverted hominoid-specific Alus show enrichment among gene regulation processes

To understand the potential contribution of inverted Alu pairs in hominoid evolution, we first identified hominoid-specific Alus using an approach inspired by Tang et al.’s human-specific mobile element detection pipeline [68]. In our approach, we used four hominoid species (chimp, gorilla, orangutan, and gibbon) and five non-hominoid species (rhesus monkey, crab-eating monkey, baboon, green monkey, and marmoset) (Figure 3A). Hominoid-specific Alus were defined as those present in at least one non-human hominoid species and absent in all non-hominoid species (see Methods). We applied a similar approach to identify hominoid-specific L1 and SVA insertions, which we provide as a resource (Supplementary Table 1). Overall, we identified 67,126 hominoid-specific Alus (Figure 3B). Most hominoid-specific Alus were from the AluY subfamily (46%; n = 30,817), followed closely by AluS (42%; n = 27,993), while a smaller fraction came from the AluJ subfamily (10%; n = 6,826), which is consistent with their evolutionary timelines [69]. Genome-wide maps aligned with previous findings on the distribution of Alu elements, with chromosome 19 being the most Alu-dense chromosome, and chromosome Y being the least (Figure 3C) [70]. Hominoid-specific Alu elements were found to be distributed relatively evenly across chromosomes, with no apparent hotspots of insertion (Figure 3C and Figure S3). Chromosome 19 has accumulated an exceptionally high number of ancestral Alus [71], which likely explains the lower ratio of hominoid-specific to total Alu elements observed on this chromosome. The notable depletion of hominoid-specific Alu elements on chromosome Y is likely attributable to its rapid evolution and high sequence divergence [72], which may hinder the identification through cross-species comparisons. At the subfamily level, we found that only hominoid-specific AluSx1 elements were significantly enriched among inverted pairs flanking skippable exons within 3,500 bp windows (adjusted *p* < 0.05, Fisher’s exact test), compared to non-hominoid elements (Figure 3D). While several other AluS and AluJ elements also appeared enriched in hominoid-specific inverted pairs, they did not attain statistical significance after multiple-testing correction. In the case of subfamily pairings, only AluSx-AluSz were enriched, reaching statistical significance at flanking windows of size 2,500 bp and 3,500 bp (adjusted *p* < 0.05).

**Figure 3.**
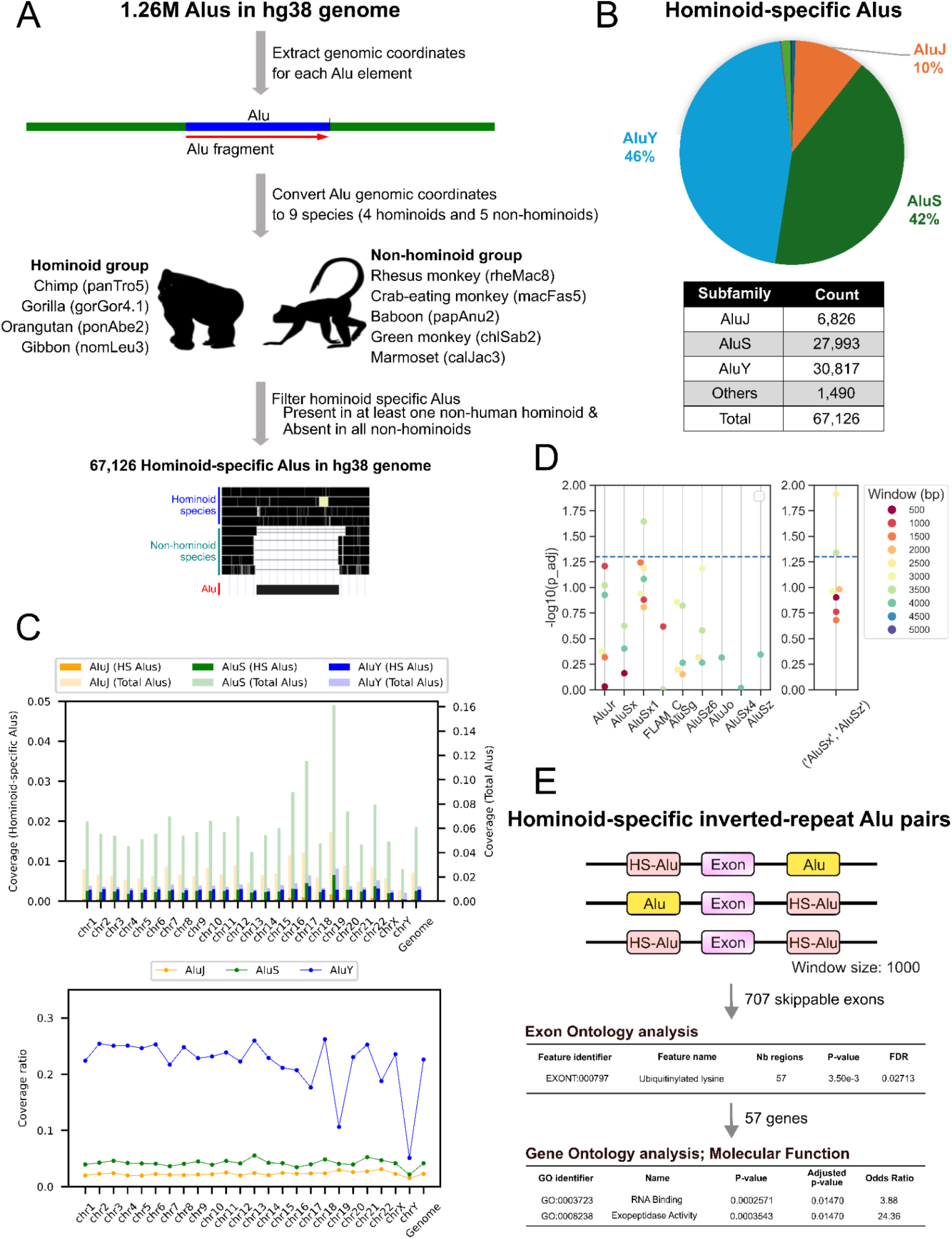
Identifying hominoid-specific Alus and their potential roles in evolution. **(A)** A comparative genomics approach to identify hominoid-specific Alus by comparing the human genome (hg38) with four hominoid species (chimp, gorilla, orangutan, and gibbon) and five non-hominoid species (rhesus monkey, crab-eating monkey, baboon, green monkey, and marmoset). **(B)** The subfamily composition of hominoid-specific Alus. **(C)** Top: The coverage of total Alus and hominoid-specific Alus across human chromosomes. Bottom: The coverage ratio (hominoid-specific Alus / total Alus) across human chromosomes. **(D)** Individual hominoid-specific Alu (left) and pairs of hominoid-specific Alu (right) subfamilies enriched in inverted orientation flanking skippable exons. Bonferroni-corrected one-tailed FE test p-values. **(E)** Exon ontology analysis and gene ontology analysis of exons flanked by hominoid-specific inverted-repeat Alu pairs.

To assess the potential roles of inverted Alus in hominoid evolution, we next performed exon ontology and gene ontology analyses. Starting from flanking windows of size 1,000 bp, we extracted skippable exons flanked by hominoid-specific inverted Alu pairs (n = 707 skippable exons). Exon ontology analysis revealed a statistically significant enrichment of “ubiquitinylated lysine” among underlying exon domains. Ubiquitinylation, more commonly referred to as ubiquitylation or ubiquitination, marks proteins for degradation and other biological processes through the addition of ubiquitin tags. Skipping these domains may lead to the removal of ubiquitination sites, potentially increasing the lifespan of the corresponding proteins and enhancing their functions. Selecting genes underlying ubiquitinated lysine (n = 57 genes), we further performed gene ontology analysis and found statistically significant enrichment of “RNA Binding” and “Exopeptidase Activity” molecular functions. Taken together, these findings suggest that inverted Alus may play a role in hominoid evolution by modulating RNA binding and exopeptidase activity through the alternative splicing of exons involved in ubiquitination-related pathways, including but not limited to protein degradation and signal transduction.

Since circRNA biogenesis has been shown to involve the formation of RNA stem loops, we further posited the relevance of hominoid-specific inverted Alu pairs in circRNA biogenesis throughout evolution. To investigate this, we quantified the enrichment ratio of circRNAs flanked by hominoid-specific inverted Alu pairs. We initially confirmed that inverted Alus lie more frequently near circRNAs compared to random genic regions of similar lengths (Figure S4A). Extending our analysis to include the hominoid specificities of underlying Alus, we found significant enrichment of inverted Alu pairs near circRNAs for all combinations of hominoid- and non-hominoid-specific Alu pairs (Figure S4B). This enrichment of inverted pairs was more pronounced in 1,000 bp windows in the absence of hominoid-specific Alus (average fold change = 2.07) compared to pairs containing a single hominoid-specific Alu (average fold change = 1.92), or both Alus being hominoid-specific (average fold change = 1.41). Given that hominoid-specific Alu elements are relatively young in evolutionary terms, we suggest that Alu pairs in which both elements are hominoid-specific may represent a baseline contribution of inverted Alu repeats to circRNA generation. It follows that the greater enrichment observed among inverted Alus lacking any hominoid specificity likely reflects an evolutionary pressure for these inverted Alu repeats to localize near circRNAs. Taken together, our findings suggest the potential selection for and widespread role of inverted Alu repeats in hominoid evolution, both in exon skipping and in other loop-involved mechanisms.

### Validation of inverted Alu pair candidates reveals hominoid-specific skipped exon isoforms

To assess whether hominoid-specific inverted Alu pairs can potentially mediate exon skipping, we selected 11 candidate exons for experimental validation. From a pool of 614 exons flanked by hominoid-specific Alu pairs within ±1000 bp windows, we initially selected 20 exons at random, spanning a range of exon inclusion levels (Supplementary Table 2). These candidates were then manually inspected in the UCSC Genome Browser, and nine were confirmed to have transcript isoforms lacking the candidate exons in the GENCODE V47 track (Figure 4A and Figure S5). We included two additional exon candidates located in *CEP290* that were not present in the original stringent skippable exon sets, but were flanked by hominoid-specific Alu pairs and had skipped isoform annotations in the UCSC Genome Browser. All 11 candidates were predicted to form RNA secondary structures containing long stem loops as compared to random genic regions (Figure 4B and Figure S6). Discriminating predicted candidate structures at nucleotide resolution revealed that the stem portion of these structures was formed by paired Alu elements (Figure 4C). We then performed RT-PCR validation for each candidate using human fibroblasts (hominoid) and African green monkey Vero cells (non-hominoid). Across all candidates, we observed the exon skipping isoform in addition to the full-length transcript in the hominoid cells only, while we solely observed the full-length transcript in non-hominoid cells (Figure 4D and Supplementary Table 3). Notably, one of our candidate genes, *CEP290*, contains two hominoid-specific inverted Alu pairs predicted to give rise to two separate skipped-exon isoforms. Excitingly, both of these skipped-exon isoforms were successfully recovered in hominoid cells and were absent in non-hominoid cells. Overall, our successful validation of all 11 putative hominoid-specific exon skipping events suggests that adjacent inverted Alu repeats likely mediate these exon skipping events during hominoid evolution.

**Figure 4.**
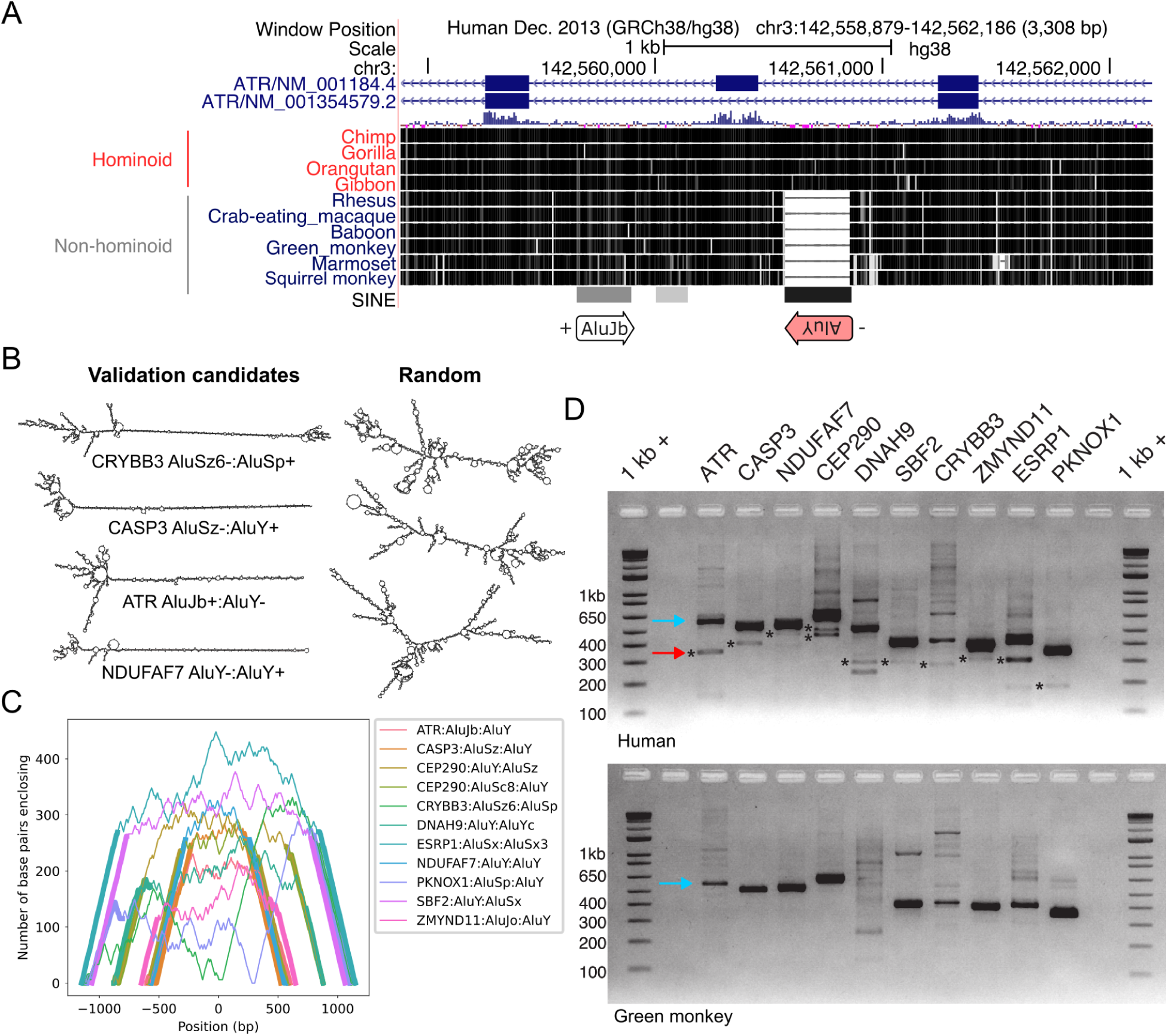
Inverted hominoid-specific Alu pairs flanking skippable exons generate splice isoforms. **(A)** UCSC Genome Browser display of a hominoid-specific skippable exon candidate that is flanked by a hominoid-specific AluY and an ancestral AluJb lying on opposite strands. **(B)** RNA secondary structure predictions of a subset of hominoid-specific validation candidates and random genic regions. **(C)** Mountain plot of all validation candidates. Bold lines indicate Alu sequences. **(D)** RT-PCR validation of hominoid-specific inverted Alu-mediated exon skipping in hominoid human HEK293T (top) compared to non-hominoid African green monkey Vero (bottom) cells. Teal arrows indicate full-length isoforms. Red arrow and stars indicate exon skipping isoforms.

## Discussion

In this work, we present an initial overview of the inverted Alu pair landscape across the human genome, and further elucidate the involvement of inverted Alu pairs in hominoid evolution. We find that inverted Alus formed by elements belonging to all major Alu lineages are generally enriched in the flanks of skippable exons at least 5,000 bp away. These inverted Alu pair sequences are predicted to form RNA stem-loops, potentially leading to exon skipping, but such inverted Alu pairs are likely not a major contributor to exon skipping overall. Rather, inverted Alu pairs may be more likely to impact exons with weakly defined 3′ acceptor splice sites. When restricting inverted Alu pairs to those found in hominoids, we uncovered their potential widespread effects impacting pathways of RNA binding and exopeptidase activity, and validated several hominoid-specific pairings suggesting the relevance of loop-out exon skipping products during evolution.

While we approached our investigation of inverted Alu pairs systematically, several key limitations should be noted. First, we assigned each Alu to its nearest exon. Considering that introns tend to be long [73], our assumption that most Alus lay close enough to form loops across only a single exon seemed reasonable. However, Alus may still be able to form pairs across multiple exons, especially in the case of short introns and exons. Indeed, such interactions have been reported [46]. Further exploration of the dynamics of Alu pairs across multiple exons, rather than the single nearest exon, is necessary.

We also emphasize that our analyses rely on the validity of underlying “skippable” and “constitutive” exon classifications. HEXEvent annotations were derived from expressed sequence tags, whereas ExonSkipDB annotations were derived from GTEx RNA-seq datasets. Neither database can truly be comprehensive in identifying skipped exon transcripts, since biological and technical factors can confound the detection of such transcripts. Alternative splicing events tend to be tissue-specific or temporal, and oftentimes cause frameshift mutations triggering nonsense-mediated mRNA decay, which render isoform detection difficult [34,74]. Therefore, improved methods with higher sensitivity that account for these challenges in skipped isoform detection are needed.

The idea that this mechanism may be more relevant for exons marked by a weak 3′ acceptor site is intriguing. Strikingly, one of the most enriched motifs among skippable exons was hnRNPA1, which is a known regulator of splicing with relevance to several cancers [75,76]. This factor is of particular interest because studies have suggested that hnRNPA1 can form homodimers that bring together the flanking regions of exons, effectively resulting in exon skipping through a loop-out mechanism [64]. Interestingly, the ability of hnRNPA1 to mediate exon skipping also relies on the presence of a weak 3′ acceptor site, and is abrogated by the introduction of a strong 3′ acceptor splice site, which could suggest the relevance of both inverted Alu pairs and hnRNPA1 in mediating a shared exon skipping mechanism. Elucidating the potential interplay between hnRNPA1 and Alu elements would benefit from follow-up studies.

While we tried to capture a variety of sequence determinants possibly involved in loop-out exon skipping, other genomic and epigenomic factors most certainly play roles in determining splicing outcomes. These factors are regulated by the cellular state and include transcription kinetics, chromatin compaction, and expression of key RNA binding proteins, which can either promote exon skipping or inclusion [4]. Further complicating these interactions are the changes to each of these components across evolution, which we are only beginning to unravel. Future work will examine how these different layers interact and change over time to regulate retrotransposon-mediated loop-out exon skipping. Our work reveals the abundance and widespread impact of inverted Alu pairs mediating exon skipping in the human genome and provides a resource of hominoid-specific retrotransposon insertions, paving the way for future exploration of the deep intronic transposon insertions and their RNA secondary structures in splicing and evolution.

## Methods

### Identifying Alu repeat pairs

We downloaded RepeatMasker annotations for repetitive elements in the human reference genome (version GRCh38.p12) and generated a BED Alu subfamily file (n=54 subfamilies) from sequences classified as “SINE_Alu” [77]. We downloaded skippable exons from two databases: HEXEvent [47] and ExonSkipDB [48]. For HEXEvent, we downloaded both cassette (skippable) and constitutive exons across all chromosomes. We selected the default parameters for cassette exons (up to an inclusion level of less than 100% of ESTs) and constitutive exons (spliced in at most 0% of ESTs) to get the full set of exons from each group to which we could later apply additional filtering. For ExonSkipDB, we selected all skippable exons derived from GTEx experiments. To remove potential confounding factors, we defined stringent mutually exclusive skippable and constitutive exon sets. Specifically, we refined the set of skippable exons by taking the union of HEXEvent and ExonSkipDB skippable exons and subtracting HEXEvent constitutive exons. Reciprocally, to generate the set of constitutive exons, we took the set of HEXEvent constitutive exons and subtracted HEXEvent and ExonSkipDB skippable exons. The number of Alu elements and exons that underlie these stringent sets are annotated in the Venn diagrams of Figure 1B.

The processing pipeline is depicted in Figure 1A. We ran BEDOPS [78] (v2.4.39) sort-bed on the RepeatMasker Alu BED file and exon BED files, followed by closest-features with the --dist flag, providing the RepeatMasker Alu BED file as file A, and each database’s exon BED file as file B. This resulted in a per-Alu annotation of each Alu element with its nearest exon, for each exon set. We grouped Alus by their nearest exons, and generated all possible combinations of Alu pairs flanking each exon. We applied a maximum flanking window cutoff of 5,000 bp measured from the start or end of each Alu element to the start or end of each exon (taking the minimum distance) with a minimum overlap of 1 bp. Inverted Alu pairs were defined as paired elements lying on opposite strands.

### Calculating enrichment of inverted repeats

We applied a flanking window cutoff ranging from 500 bp to 5,000 bp at 500 bp increments, taken from each side of the exon, which we measured from the start or end of each Alu element to the start or end of each exon (taking the minimum distance). We used a minimum overlap of 1 bp therefore allowing Alu elements to fall partially outside of flanking windows. We constructed 2 by 2 contingency tables with columns denoting exon types: skippable and constitutive, and rows denoting Alu pairing types: inverted and non-inverted. For hominoid-specific Alus, we instead used rows indicating the hominoid specificities within inverted Alu pairs (hominoid-specific and non-specific). In the case of Alu subfamily enrichment, we further restricted rows to include at least one member (Figure 1F and Figure 3D left) or both members (Figure 1F and Figure 3D right) of the pair belonging to the specified Alu subfamily. We calculated p-values using a one-tailed Fisher’s exact test and applied family-wise Bonferroni multiple-testing correction.

### Evaluating the importance of various exon and Alu features

We compared upstream and downstream Alu lengths, upstream and downstream Alu distances to their flanked exon, intra-Alu distances, and exon lengths across exon type and Alu inversion combinations. To account for overpowered comparisons, we adopted a 10-fold randomized sampling of 1,000 pairs within each category, and compared categories using scipy’s two-sample two-sided Kolmogorov–Smirnov (KS) test [79] while adjusting p-values by applying the Bonferroni method.

### Predicting RNA secondary structure and minimum free energy

We extracted strand-specific RNA sequences for Alu pairs (referred to as Alu 1 and Alu 2) spanning the start of Alu 1, through the intervening flanked exon, and extending until the end of Alu 2 using bedtools getfasta -s with human reference genome hg38 (Figure 2A). We split the resulting FASTA file into files of 100 sequences each using split --suffix-length=6 -d −l 200 to facilitate parallel processing. We ran RNAfold [55,56] (v2.5.0) using default parameters on each file and extracted predicted MFE values for each sequence. We generated a set of size-matched random genic sequences for each exon type and Alu pair inversion sequence set for an appropriate baseline. In other words, for every Alu1-exon-Alu2 sequence in our sets of interest, we chose a region of the same length within refGene gene regions [80] (vGCA_000001405.15) using bedtools shuffle -incl <refGene_TXT> -g <hg38_FASTA> -i <exon_type_alu_inversion_BED>. For 1 kb windows, RNAfold processing of all sequences of interest completed successfully, whereas for 2 kb and 5 kb windows, a small fraction (0.13%-0.21%) of sequence runs failed and did not produce MFE values. Likewise, for the random sets of sequences, a small fraction (0.24%) of sequence runs failed as well. To account for the increased structural stability observed for longer RNA sequences, we scaled MFE scores according to sequence length (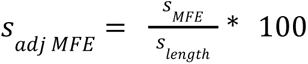) [81]. Since these experiments were overpowered and all p-values were extremely significant, we chose to report effect sizes as measured by Cohen’s *d* instead.

### Assigning tissue-specific PSI

We downloaded percent spliced-in annotations for human exons from the ASCOT alternative splicing catalog which cover 53 GTEx tissues and 3 additional retinal tissues [59]. When plotting the minimum tissue PSI, we excluded exons with a minimum PSI < 10% across all 56 tissues and those with a minimum tissue PSI > 90% to remove exons at the extremes of exclusion and inclusion within particular tissues.

### Defining exon strength

We computed exon definition scores for both 3′ acceptor and 5′ donor splice sites using the maxentpy Python library implementation of MaxEntScan [82,83]. We labeled exons according to their 3′ acceptor splice site score followed by their 5′ donor splice site score, where a score greater than 8 was defined as strong (S) and a score < 6 was defined as weak (W) in line with previous studies [57].

### RNA editing burden

We downloaded hg38 RNA editing sites from REDIportal [84] which we converted to a BED file. We overlapped editing sites with Alu pair sequences in Python using pybedtools [85].

### Motif enrichment analysis

We performed differential motif enrichment analysis using the Docker image of SEA [86] (v5.5.5) on sequences of inverted Alu pairs flanking skippable exons defined by weak 3′ acceptor and strong 5′ donor (WS) splice sites. Sequences containing exons with a median PSI < 50% across all tissues in ASCOT were used as primary sequences, whereas those with a median PSI > 50% across tissues were used as control sequences (x-axis of Figure 2F). Enriched motifs were compared to those discovered in median PSI > 50% primary sequences using median PSI < 50% sequences as controls (y-axis). P-values were calculated by SEA using Fisher’s exact test and adjusted using the Benjamini–Hochberg procedure. When multiple motifs were reported for the same RNA binding protein, p-values were combined using Fisher’s method. Known motifs were curated from the ATtRACT RNA binding protein motif database [87] by selecting position weight matrices from *Homo sapiens* in which the target gene was not mutated. We used the chen2meme utility from the MEME Suite [88] to convert the motif database file to a format that is compatible with MEME Suite tools.

### Identifying hominoid-specific transposable elements

We identified hominoid-specific transposable elements using an approach inspired by Tang et al.’s human-specific mobile element detection pipeline [68]. We extracted the genomic coordinates of each transposable element’s insertion site annotated in the human reference genome (version GRCh38.p12), mainly focusing on retrotransposons including LINEs, Alus, and SVAs. We then compared these genomic regions to the orthologous regions of the genomes of four hominoid species and six non-hominoid primates, using the web version of UCSC LiftOver [89]. We opted to apply lenient selection criteria to prioritize finding as many hominoid-specific transposable elements as possible, rather than potentially filtering out candidates due to limitations in existing annotations. Transposable elements were considered to be hominoid-specific if their insertion sites were present in at least three hominoids but were absent in all non-hominoid primates. We assigned the hominoid group to include human (hg38), chimp (panTro6), gorilla (gorGor6), orangutan (ponAbe3), and gibbon (nomLeu3). The non-hominoid group consisted of rhesus (rheMac10), crab-eating macaque (macFas5), baboon (papAnu4), green monkey (chlSab2), marmoset (calJac4), and squirrel monkey (saiBol1). We defined hominoid-specific Alu pairs as those pairs that include at least one hominoid-specific Alu element.

### Calculating Alu density across human chromosomes

To determine the total and hominoid-specific Alu density across human chromosomes, we calculated the length of each Alu element and summed these lengths to obtain the total length of Alu sequences per chromosome. We then calculated the Alu density by dividing the summed Alu length by the corresponding chromosome length. The coverage ratio was subsequently obtained by dividing the hominoid-specific Alu density by the total Alu density.

### Exon ontology analysis and GO Molecular Function analysis

Exon ontology analysis was performed on the set of skippable exons flanked by hominoid-specific inverted-repeat Alu pairs using the Exon Ontology database [90]. As Exon Ontology relies on the hg19 genome build, genomic coordinates were converted to human hg19 using the web version of UCSC LiftOver. Genes containing skippable exons that were included in the exon ontology feature with statistical significance were used to perform Gene Ontology Molecular Function analysis using the web-based tool, EnrichR [90–93]. Significant results for the exon ontology analysis and the gene ontology analysis were reported (Benjamini-Hochberg adjusted p-value < 0.05).

### Selecting and validating candidate inverted Alu pairs

We selected twenty putative inverted Alu-mediated exon skipping events, spanning a broad range of exon inclusion levels (50-99%) as reported in ASCOT [59]. Each event was manually verified using the UCSC Genome Browser to confirm the presence of at least one hominoid-specific Alu element flanking the skippable exon and the presence of transcript isoforms lacking the candidate exons in the GENCODE V47 track (Figure 4A and Figure S5). From these, we selected nine exons for further experimental validation. We also included two additional exon candidates located in *CEP290* that were not present in the original stringent skippable exon sets, but were flanked by hominoid-specific Alu pairs and had associated skipped isoform track annotations. In total, ten genes encompassing 11 exon skipping events passed the selection criteria for experimental validation (Supplementary Table 2).

Approximately 1×10^5 human fibroblasts and Vero cells (CCL-81, adult African green monkey) were seeded in 12-well plates with DMEM medium (Gibco Laboratories, Gaithersburg, MD, USA) supplemented with 10% fetal bovine serum (Gibco) and 1X MEM Non-Essential Amino Acids Solution (Thermo Fisher Scientific). To comprehensively capture alternative splicing isoforms, cells were treated with the nonsense-mediated decay inhibitor cycloheximide (CHX) at 100 ug/mL for 6 hours prior to RNA extraction. Total RNA was isolated using the PureLink RNA Mini Kit (Invitrogen), with concentration and quality assessed using a NanoDrop OneC (Thermo Fisher Scientific). cDNA was generated via reverse transcription with SuperScript IV VILO Mastermix (Thermo Fisher Scientific) following the manufacturer’s protocol (25°C for 10 min, 50°C for 10 min, 85°C for 5 min).

RT-PCR validation of inverted Alu-mediated exon skipping was performed using KAPA2G Robust HotStart ReadyMix (KK5702; KAPA Biosystems). The cycling program consisted of: 95°C for 5 min; 35 cycles of 95°C for 30 sec, 60°C for 30 sec, and 72°C for 30 sec; a final 72°C extension for 30 sec; and hold at 4°C. RT-PCR products were analyzed by 2% agarose gel electrophoresis. Exon skipping bands with the expected size were excised, extracted (PureLink Quick Gel Extraction, Invitrogen), and confirmed by either Sanger sequencing or Amplicon-EZ (Azenta, Inc.). Validation primer sequences can be found in Supplementary Table 3.

### Enrichment of inverted Alus flanking circRNA

We tested for the enrichment of inverted Alu pairs within the flanks of circRNA using known human circRNAs [94], reference Alus [77], and hominoid-specific Alus that we identified in this study. Intergenic regions were extracted and randomized to match the length distribution of the circRNA annotations, using the shuffling method implemented in regioneR [95]. We assigned each Alu element to its closest randomized region located either upstream or downstream. To identify regions containing inverted Alus, we selected regions containing Alu elements in both their upstream and downstream flanks lying on opposite strands within windows of size 1,000 bp, 2,000 bp, and 5,000 bp. This process was repeated for 10,000 iterations to ensure robust randomization. The number of randomized regions containing inverted Alu pairs was used for analyses.

## Abbreviations

circRNA: circular RNA
W: weak
S: strong
SS: strong 3′ acceptor splice site, strong 5′ donor splice site
SW: strong 3′ acceptor splice site, weak 5′ donor splice site
WS: weak 3′ acceptor splice site, strong 5′ donor splice site
WW: weak 3′ acceptor splice site, weak 5′ donor splice site
PSI: percent spliced-in

## Declarations

## Ethics approval and consent to participate

Not applicable

## Consent for publication

Not applicable

## Availability of data and materials

RepeatMasker Alu annotations were downloaded from the UCSC Genome Browser. Exons were downloaded from HEXEvent and ExonSkipDB. We performed our analyses in Python (v3.9.18). Jupyter notebooks, supporting scripts, and environment files are available on GitHub at https://github.com/dcdxy/inverted_alus.

## Authors’ contributions

B.Z. and E.A.L. conceived the project, and D.D., J. Kim., B.Z., and E.A.L. designed the analyses and experiments. D.D. conducted genome-wide Alu characterization and secondary structure prediction analyses. J. Kim and B.Z. performed hominoid-specific Alu analyses and PCR validation. J. Ku and J. Kim carried out circRNA analyses. D.D. wrote the manuscript with contributions from all authors. B.Z. and E.A.L. supervised the study and revised the manuscript. All authors read and approved the final manuscript.

## Acknowledgements

We thank Junseok Park for statistical and computing advice, and members of the Lee lab for helpful discussions.

## References

1. Merkin J, Russell C, Chen P, Burge CB. Evolutionary dynamics of gene and isoform regulation in Mammalian tissues. Science. 2012 Dec 21;338(6114):1593–9.

2. Baralle FE, Giudice J. Alternative splicing as a regulator of development and tissue identity. Nat Rev Mol Cell Biol. 2017 Jul;18(7):437–51.

3. Mazin PV, Khaitovich P, Cardoso-Moreira M, Kaessmann H. Alternative splicing during mammalian organ development. Nat Genet. 2021 Jun;53(6):925–34.

4. Marasco LE, Kornblihtt AR. The physiology of alternative splicing. Nat Rev Mol Cell Biol. 2023 Apr;24(4):242–54.

5. Wang Z, Burge CB. Splicing regulation: from a parts list of regulatory elements to an integrated splicing code. RNA. 2008 May;14(5):802–13.

6. Abramowicz A, Gos M. Splicing mutations in human genetic disorders: examples, detection, and confirmation. J Appl Genet. 2018 Aug;59(3):253–68.

7. Ule J, Blencowe BJ. Alternative splicing regulatory networks: Functions, mechanisms, and evolution. Mol Cell. 2019 Oct 17;76(2):329–45.

8. Liu S, Cheng C. Alternative RNA splicing and cancer. Wiley Interdiscip Rev RNA. 2013 Sep;4(5):547–66.

9. Keren H, Lev-Maor G, Ast G. Alternative splicing and evolution: diversification, exon definition and function. Nat Rev Genet. 2010 May;11(5):345–55.

10. Kim E, Magen A, Ast G. Different levels of alternative splicing among eukaryotes. Nucleic Acids Res. 2007;35(1):125–31.

11. Barbosa-Morais NL, Irimia M, Pan Q, Xiong HY, Gueroussov S, Lee LJ, et al. The evolutionary landscape of alternative splicing in vertebrate species. Science. 2012 Dec 21;338(6114):1587–93.

12. Reyes A, Anders S, Weatheritt RJ, Gibson TJ, Steinmetz LM, Huber W. Drift and conservation of differential exon usage across tissues in primate species. Proc Natl Acad Sci U S A. 2013 Sep 17;110(38):15377–82.

13. Calarco JA, Xing Y, Cáceres M, Calarco JP, Xiao X, Pan Q, et al. Global analysis of alternative splicing differences between humans and chimpanzees. Genes Dev. 2007 Nov 15;21(22):2963–75.

14. Hiller M, Zhang Z, Backofen R, Stamm S. Pre-mRNA secondary structures influence exon recognition. PLoS Genet. 2007 Nov;3(11):e204.

15. Cao X, Zhang Y, Ding Y, Wan Y. Identification of RNA structures and their roles in RNA functions. Nat Rev Mol Cell Biol. 2024 Jun 26;25(10):784–801.

16. Shepard PJ, Hertel KJ. Conserved RNA secondary structures promote alternative splicing. RNA. 2008 Aug;14(8):1463–9.

17. Bartys N, Kierzek R, Lisowiec-Wachnicka J. The regulation properties of RNA secondary structure in alternative splicing. Biochim Biophys Acta Gene Regul Mech. 2019 Nov;1862(11-12):194401.

18. Solnick D. Alternative splicing caused by RNA secondary structure. Cell. 1985 Dec;43(3 Pt 2):667–76.

19. Eperon LP, Graham IR, Griffiths AD, Eperon IC. Effects of RNA secondary structure on alternative splicing of pre-mRNA: is folding limited to a region behind the transcribing RNA polymerase? Cell. 1988 Jul 29;54(3):393–401.

20. Cao D. Reverse complementary matches simultaneously promote both back-splicing and exon-skipping. BMC Genomics. 2021 Aug 3;22(1):586.

21. Kim Y, Park J, Kim S, Kim M, Kang MG, Kwak C, et al. PKR senses nuclear and mitochondrial signals by interacting with endogenous double-stranded RNAs. Mol Cell. 2018 Sep 20;71(6):1051–63.e6.

22. Cost GJ, Feng Q, Jacquier A, Boeke JD. Human L1 element target-primed reverse transcription in vitro. EMBO J. 2002 Nov 1;21(21):5899–910.

23. Kojima KK. Human transposable elements in Repbase: genomic footprints from fish to humans. Mob DNA. 2018 Jan 4;9:2.

24. Chen LL, Yang L. ALUternative regulation for gene expression. Trends Cell Biol. 2017 Jul;27(7):480–90.

25. Häsler J, Samuelsson T, Strub K. Useful “junk”: Alu RNAs in the human transcriptome. Cell Mol Life Sci. 2007 Jul;64(14):1793–800.

26. Konkel MK, Walker JA, Batzer MA. LINEs and SINEs of primate evolution. Evol Anthropol. 2010 Nov 1;19(6):236–49.

27. Mighell AJ, Markham AF, Robinson PA. Alu sequences. FEBS Lett. 1997 Nov 3;417(1):1–5.

28. Batzer MA, Deininger PL, Hellmann-Blumberg U, Jurka J, Labuda D, Rubin CM, et al. Standardized nomenclature for Alu repeats. J Mol Evol. 1996 Jan;42(1):3–6.

29. Szitenberg A, Cha S, Opperman CH, Bird DM, Blaxter ML, Lunt DH. Genetic drift, not life history or RNAi, determine long-term evolution of transposable elements. Genome Biol Evol. 2016 Oct 5;8(9):2964–78.

30. Catlin NS, Josephs EB. The important contribution of transposable elements to phenotypic variation and evolution. Curr Opin Plant Biol. 2022 Feb;65(102140):102140.

31. Bourque G, Burns KH, Gehring M, Gorbunova V, Seluanov A, Hammell M, et al. Ten things you should know about transposable elements. Genome Biol. 2018 Nov 19;19(1):199.

32. Batzer MA, Deininger PL. Alu repeats and human genomic diversity. Nat Rev Genet. 2002 May;3(5):370–9.

33. Deininger P. Alu elements: know the SINEs. Genome BiolGenome Biology. 2011;12(12):236.

34. Sibley CR, Blazquez L, Ule J. Lessons from non-canonical splicing. Nat Rev Genet. 2016 Jul;17(7):407–21.

35. Lander ES, Linton LM, Birren B, Nusbaum C, Zody MC, Baldwin J, et al. Initial sequencing and analysis of the human genome. Nature. 2001 Feb 15;409(6822):860–921.

36. Kim DDY, Kim TTY, Walsh T, Kobayashi Y, Matise TC, Buyske S, et al. Widespread RNA Editing of Embedded Alu Elements in the Human Transcriptome. Genome Res. 2004 Sep 1;14(9):1719–25.

37. Cao X, Zhang Y, Payer LM, Lords H, Steranka JP, Burns KH, et al. Polymorphic mobile element insertions contribute to gene expression and alternative splicing in human tissues. Genome Biol. 2020 Jul 27;21(1):185.

38. Konkel MK, Walker JA, Hotard AB, Ranck MC, Fontenot CC, Storer J, et al. Sequence analysis and characterization of active human Alu subfamilies based on the 1000 genomes pilot project. Genome Biol Evol. 2015 Aug 29;7(9):2608–22.

39. Ganguly A, Dunbar T, Chen P, Godmilow L, Ganguly T. Exon skipping caused by an intronic insertion of a young Alu Yb9 element leads to severe hemophilia A. Hum Genet. 2003 Sep;113(4):348–52.

40. Claverie-Martín F, Flores C, Antón-Gamero M, González-Acosta H, García-Nieto V. The Alu insertion in the CLCN5 gene of a patient with Dent’s disease leads to exon 11 skipping. J Hum Genet. 2005 Jul 23;50(7):370–4.

41. Wallace MR, Andersen LB, Saulino AM, Gregory PE, Glover TW, Collins FS. A de novo Alu insertion results in neurofibromatosis type 1. Nature. 1991 Oct 31;353(6347):864–6.

42. Zhao B, Nguyen MA, Woo S, Kim J, Yu TW, Lee EA. Contribution and therapeutic implications of retroelement insertions in ataxia telangiectasia. Am J Hum Genet. 2023 Nov 2;110(11):1976–82.

43. Zhang XO, Wang HB, Zhang Y, Lu X, Chen LL, Yang L. Complementary sequence-mediated exon circularization. Cell. 2014 Sep 25;159(1):134–47.

44. Lee K, Ku J, Ku D, Kim Y. Inverted Alu repeats: friends or foes in the human transcriptome. Exp Mol Med. 2024 Jun 14;56(6):1250–62.

45. Masson E, Maestri S, Bordeau V, Cooper DN, Férec C, Chen JM. Alu insertion-mediated dsRNA structure formation with pre-existing Alu elements as a disease-causing mechanism. Am J Hum Genet. 2024 Sep 4;111(10):2176–89.

46. Xia B, Zhang W, Zhao G, Zhang X, Bai J, Brosh R, et al. On the genetic basis of tail-loss evolution in humans and apes. Nature. 2024 Feb 29;626(8001):1042–8.

47. Busch A, Hertel KJ. HEXEvent: a database of Human EXon splicing Events. Nucleic Acids Res. 2013 Jan;41(Database issue):D118–24.

48. Kim P, Yang M, Yiya K, Zhao W, Zhou X. ExonSkipDB: functional annotation of exon skipping event in human. Nucleic Acids Res. 2020 Jan 8;48(D1):D896–907.

49. Cook GW, Konkel MK, Major JD 3rd, Walker JA, Han K, Batzer MA. Alu pair exclusions in the human genome. Mob DNA. 2011 Sep 23;2(1):10.

50. Payer LM, Steranka JP, Kryatova MS, Grillo G, Lupien M, Rocha PP, et al. Alu insertion variants alter gene transcript levels. Genome Res. 2021 Dec;31(12):2236–48.

51. Lev-Maor G, Ram O, Kim E, Sela N, Goren A, Levanon EY, et al. Intronic Alus influence alternative splicing. PLoS Genet. 2008 Sep 26;4(9):e1000204.

52. Payer LM, Steranka JP, Ardeljan D, Walker J, Fitzgerald KC, Calabresi PA, et al. Alu insertion variants alter mRNA splicing. Nucleic Acids Res. 2019 Jan 10;47(1):421–31.

53. Cui Y, Cai M, Stanley HE. Comparative analysis and classification of cassette exons and constitutive exons. Biomed Res Int. 2017 Dec 4;2017:7323508.

54. Zheng CL, Fu XD, Gribskov M. Characteristics and regulatory elements defining constitutive splicing and different modes of alternative splicing in human and mouse. RNA. 2005 Dec;11(12):1777–87.

55. Lorenz R, Bernhart SH, Höner Zu Siederdissen C, Tafer H, Flamm C, Stadler PF, et al. ViennaRNA Package 2.0. Algorithms Mol Biol. 2011 Nov 24;6:26.

56. Gruber AR, Lorenz R, Bernhart SH, Neuböck R, Hofacker IL. The Vienna RNA websuite. Nucleic Acids Res. 2008 Jul 1;36(Web Server issue):W70–4.

57. Movassat M, Forouzmand E, Reese F, Hertel KJ. Exon size and sequence conservation improves identification of splice-altering nucleotides. RNA. 2019 Dec;25(12):1793–805.

58. Shepard PJ, Choi EA, Busch A, Hertel KJ. Efficient internal exon recognition depends on near equal contributions from the 3’ and 5’ splice sites. Nucleic Acids Res. 2011 Nov 1;39(20):8928–37.

59. Ling JP, Wilks C, Charles R, Leavey PJ, Ghosh D, Jiang L, et al. ASCOT identifies key regulators of neuronal subtype-specific splicing. Nat Commun. 2020 Jan 9;11(1):137.

60. Nishikura K. A-to-I editing of coding and non-coding RNAs by ADARs. Nat Rev Mol Cell Biol. 2016 Feb;17(2):83–96.

61. Zhu W, Zhou BL, Rong LJ, Ye L, Xu HJ, Zhou Y, et al. Roles of PTBP1 in alternative splicing, glycolysis, and oncogensis. J Zhejiang Univ Sci B. 2020 Feb;21(2):122–36.

62. Huang S, Luo K, Jiang L, Zhang XD, Lv YH, Li RF. PCBP1 regulates the transcription and alternative splicing of metastasis-related genes and pathways in hepatocellular carcinoma. Sci Rep. 2021 Dec 2;11(1):23356.

63. Martinez FJ, Pratt GA, Van Nostrand EL, Batra R, Huelga SC, Kapeli K, et al. Protein-RNA networks regulated by normal and ALS-associated mutant HNRNPA2B1 in the nervous system. Neuron. 2016 Nov 23;92(4):780–95.

64. Blanchette M, Chabot B. Modulation of exon skipping by high-affinity hnRNP A1-binding sites and by intron elements that repress splice site utilization. EMBO J. 1999 Apr 1;18(7):1939–52–1952.

65. Shao M, Hao S, Jiang L, Cai Y, Zhao X, Chen Q, et al. CRIT: Identifying RNA-binding protein regulator in circRNA life cycle via non-negative matrix factorization. Mol Ther Nucleic Acids. 2022 Dec 13;30:398–406.

66. Huang C, Yang Y, Wang X, Chen S, Liu Z, Li Z, et al. PTBP1-mediated biogenesis of circATIC promotes progression and cisplatin resistance of bladder cancer. Int J Biol Sci. 2024 Jun 24;20(9):3570–89.

67. Amini J, Zafarjafarzadeh N, Ghahramanlu S, Mohammadalizadeh O, Mozaffari E, Bibak B, et al. Role of Circular RNA MMP9 in Glioblastoma Progression: From Interaction With hnRNPC and hnRNPA1 to Affecting the Expression of BIRC5 by Sequestering miR-149. J Mol Recognit. 2024 Oct 14;e3109.

68. Tang W, Mun S, Joshi A, Han K, Liang P. Mobile elements contribute to the uniqueness of human genome with 15,000 human-specific insertions and 14 Mbp sequence increase. DNA Res. 2018 Oct 1;25(5):521–33.

69. Dagan T, Sorek R, Sharon E, Ast G, Graur D. AluGene: a database of Alu elements incorporated within protein-coding genes. Nucleic Acids Res. 2004 Jan 1;32(Database issue):D489–92.

70. Grover D, Mukerji M, Bhatnagar P, Kannan K, Brahmachari SK. Alu repeat analysis in the complete human genome: trends and variations with respect to genomic composition. Bioinformatics. 2004 Apr 12;20(6):813–7.

71. Grimwood J, Gordon LA, Olsen A, Terry A, Schmutz J, Lamerdin J, et al. The DNA sequence and biology of human chromosome 19. Nature. 2004 Apr 1;428(6982):529–35.

72. Kuderna LFK, Ulirsch JC, Rashid S, Ameen M, Sundaram L, Hickey G, et al. Identification of constrained sequence elements across 239 primate genomes. Nature. 2024 Jan;625(7996):735–42.

73. Fox-Walsh KL, Dou Y, Lam BJ, Hung SP, Baldi PF, Hertel KJ. The architecture of pre-mRNAs affects mechanisms of splice-site pairing. Proc Natl Acad Sci U S A. 2005 Nov 8;102(45):16176–81.

74. Pan Q, Shai O, Lee LJ, Frey BJ, Blencowe BJ. Deep surveying of alternative splicing complexity in the human transcriptome by high-throughput sequencing. Nat Genet. 2008 Dec;40(12):1413–5.

75. Babic I, Anderson ES, Tanaka K, Guo D, Masui K, Li B, et al. EGFR mutation-induced alternative splicing of max contributes to growth of glycolytic tumors in brain cancer. Cell Metab. 2013 Jun;17(6):1000–8.

76. Zhang Y, Qian J, Gu C, Yang Y. Alternative splicing and cancer: a systematic review. Signal Transduct Target Ther. 2021 Feb 24;6(1):78.

77. Smit AFA, Hubley R, Green P. RepeatMasker Open-4.0 [Internet]. 2013-2015. Available from: http://www.repeatmasker.org

78. Neph S, Kuehn MS, Reynolds AP, Haugen E, Thurman RE, Johnson AK, et al. BEDOPS: high-performance genomic feature operations. Bioinformatics. 2012 Jul 15;28(14):1919–20.

79. Virtanen P, Gommers R, Oliphant TE, Haberland M, Reddy T, Cournapeau D, et al. SciPy 1.0: fundamental algorithms for scientific computing in Python. Nat Methods. 2020 Mar;17(3):261–72.

80. Pruitt KD, Tatusova T, Maglott DR. NCBI reference sequences (RefSeq): a curated non-redundant sequence database of genomes, transcripts and proteins. Nucleic Acids Res. 2007 Jan;35(Database issue):D61–5.

81. Zhang BH, Pan XP, Cox SB, Cobb GP, Anderson TA. Evidence that miRNAs are different from other RNAs. Cell Mol Life Sci. 2006 Jan;63(2):246–54.

82. Yeo G, Burge CB. Maximum entropy modeling of short sequence motifs with applications to RNA splicing signals. J Comput Biol. 2004;11(2-3):377–94.

83. Zhang XO. maxentpy [Internet]. 2022. Available from: https://github.com/kepbod/maxentpy

84. Picardi E, D’Erchia AM, Lo Giudice C, Pesole G. REDIportal: a comprehensive database of A-to-I RNA editing events in humans. Nucleic Acids Res. 2017 Jan 4;45(D1):D750–7.

85. Dale RK, Pedersen BS, Quinlan AR. Pybedtools: a flexible Python library for manipulating genomic datasets and annotations. Bioinformatics. 2011 Dec 15;27(24):3423–4.

86. Bailey TL, Grant CE. SEA: Simple Enrichment Analysis of motifs [Internet]. bioRxiv. 2021. Available from: 10.1101/2021.08.23.457422

87. Giudice G, Sánchez-Cabo F, Torroja C, Lara-Pezzi E. ATtRACT-a database of RNA-binding proteins and associated motifs. Database (Oxford). 2016 Apr 7;2016:baw035.

88. Bailey TL, Johnson J, Grant CE, Noble WS. The MEME suite. Nucleic Acids Res. 2015 Jul 1;43(W1):W39–49.

89. Karolchik D, Hinrichs AS, Furey TS, Roskin KM, Sugnet CW, Haussler D, et al. The UCSC Table Browser data retrieval tool. Nucleic Acids Res. 2004 Jan 1;32(Database issue):D493–6.

90. Tranchevent LC, Aubé F, Dulaurier L, Benoit-Pilven C, Rey A, Poret A, et al. Identification of protein features encoded by alternative exons using Exon Ontology. Genome Res. 2017 Jun;27(6):1087–97.

91. Chen EY, Tan CM, Kou Y, Duan Q, Wang Z, Meirelles GV, et al. Enrichr: interactive and collaborative HTML5 gene list enrichment analysis tool. BMC Bioinformatics. 2013 Apr 15;14(1):128.

92. Kuleshov MV, Jones MR, Rouillard AD, Fernandez NF, Duan Q, Wang Z, et al. Enrichr: a comprehensive gene set enrichment analysis web server 2016 update. Nucleic Acids Res. 2016 Jul 8;44(W1):W90–7.

93. Xie Z, Bailey A, Kuleshov MV, Clarke DJB, Evangelista JE, Jenkins SL, et al. Gene set knowledge discovery with Enrichr. Curr Protoc. 2021 Mar;1(3):e90.

94. Wu W, Ji P, Zhao F. CircAtlas: an integrated resource of one million highly accurate circular RNAs from 1070 vertebrate transcriptomes. Genome Biol. 2020 Apr 28;21(1):101.

95. Gel B, Díez-Villanueva A, Serra E, Buschbeck M, Peinado MA, Malinverni R. regioneR: an R/Bioconductor package for the association analysis of genomic regions based on permutation tests. Bioinformatics. 2016 Jan 15;32(2):289–91.

