## Supplemental Figures for "Inverted Alu repeats in loop-out exon skipping across hominoid evolution"

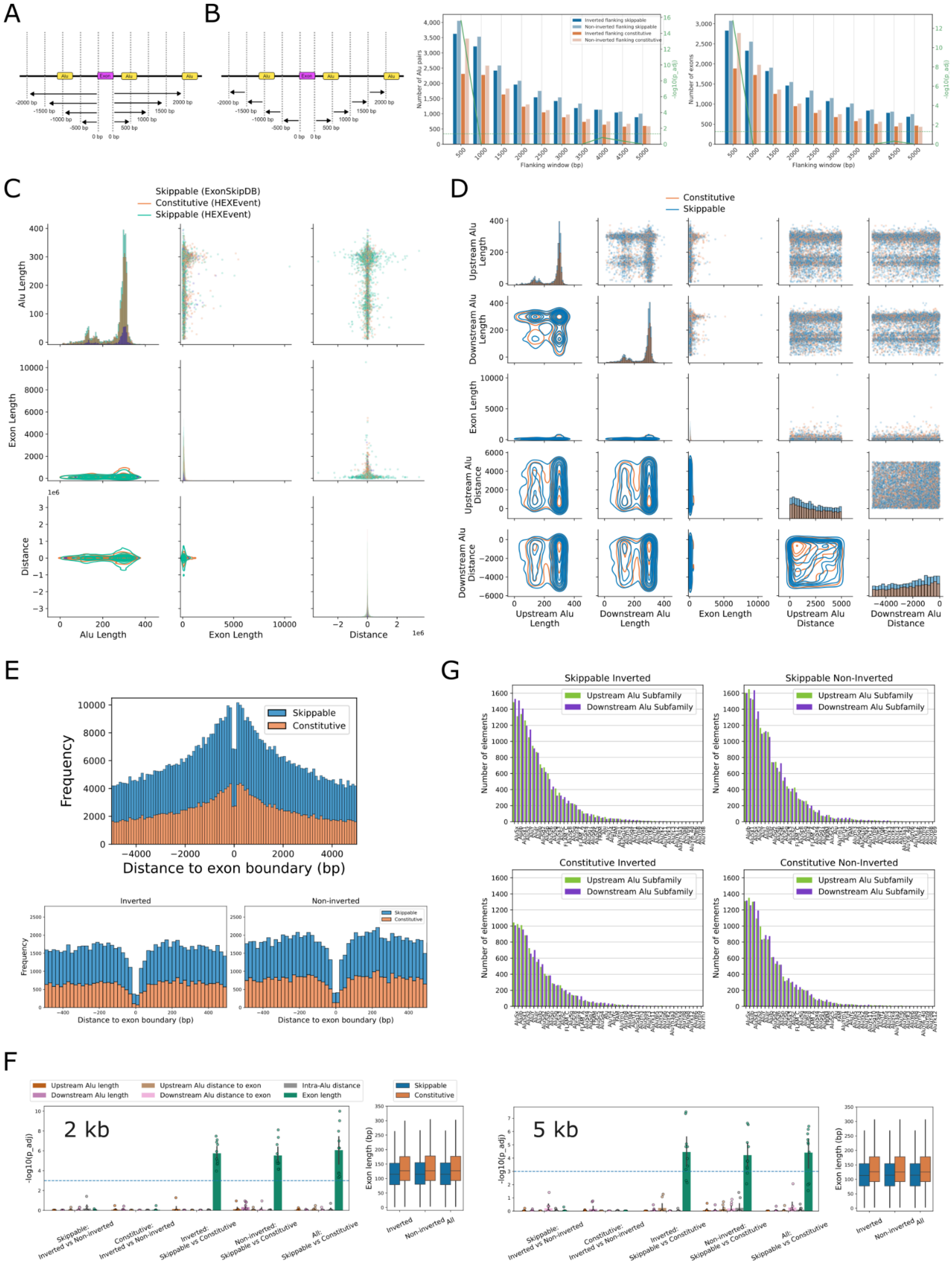
A
B
Flanking window (bp)
Number of Alu pairs
Number of exons
-log10(p_adj)
C
Skippable (ExonSkipDB)
Constitutive (HEXEvent)
Skippable (HEXEvent)
Alu Length
Exon Length
Distance
D
Constitutive
Skippable
Upstream Alu Length
Downstream Alu Length
Exon Length
Upstream Alu Distance
Downstream Alu Distance
E
Skippable
Constitutive
Frequency
Distance to exon boundary (bp)
Inverted
Non-inverted
G
Skippable Inverted
Skippable Non-Inverted
Constitutive Inverted
Constitutive Non-Inverted
Upstream Alu Subfamily
Downstream Alu Subfamily
Number of elements
F
Upstream Alu length
Downstream Alu length
Upstream Alu distance to exon
Downstream Alu distance to exon
Intra-Alu distance
Exon length
Skippable
Constitutive
2 kb
5 kb
-log10(p_adj)
Exon length (bp)
Skippable: Inverted vs Non-inverted
Constitutive: Inverted vs Non-inverted
Inverted: Skippable vs Constitutive
Non-inverted: Skippable vs Constitutive
All: Skippable vs Constitutive
Inverted
Non-inverted
All

**Figure S1.** Characterizing inverted Alus genome-wide.

**(A)** Windows are defined as overlapping intervals spanning from either end of the exon to the shortest distance to each Alu, i.e. the end of the upstream Alu or the start of the downstream Alu. In this schematic, the innermost Alu pair flanking the exon falls under windows of size {1000 bp, 1500 bp, 2000 bp, …, 5000 bp}. The outermost Alu pair flanking the exon falls under windows of size {2000 bp, 2500 bp, 3000 bp, …, 5000 bp}.

**(B)** Schematic (left) depicts a non-overlapping scheme of windows used to calculate symmetric inverted Alu pair enrichment. As in (A), the shortest distances between Alu and exon boundaries are used to determine window inclusion. In this example, neither pair falls into any symmetric window. Bar plots show the number of Alu pairs (middle) and exons (right) satisfying symmetric window conditions. Overlaid green lines indicate -log10 of Bonferroni-adjusted one-tailed Fisher's exact test p-values.

**(C)** Distributions of Alu length, nearest exon length, and distance in bp to nearest exon of all fixed Alus. Random sampling of 10,000 Alus.

**(D)** Distributions of upstream and downstream Alu lengths, exon lengths, and distance in bp to exon of upstream and downstream Alus of Alu pairs falling within +/- 5000 bp windows.

**(E)** Top: Distance to exon boundary of Alus within non-inverted pairings. Bottom: Zoomed-in view of the central 500 bp surrounding exon boundaries across inverted (left) and non-inverted (right) pairs.

**(F**) Enrichment of various length and distance features across Alu pair inversion and exon type. Significance is calculated from 10-fold random sampling of 1000 pairs of Alus within 2000 bp (left) or 5000 bp (right) flanking windows. Bonferroni-corrected Kolmogorov–Smirnov test p-values.

**(G)** Frequency of each Alu subfamily in +/- 5000 bp windows.

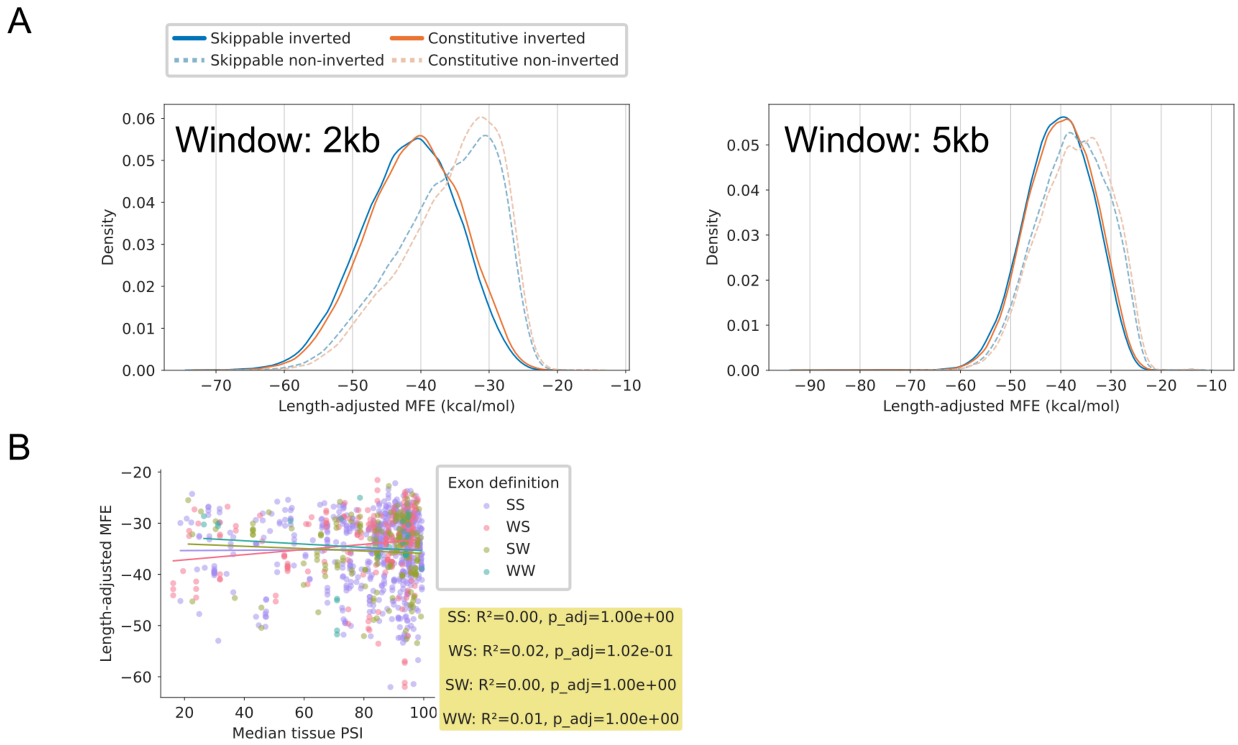


**Figure S2.** Predicting RNA secondary structure across different window sizes.

**(A)** MFE distributions across exon type and Alu inversion categories using flanking windows of size 2000 bp (left) and 5000 bp (right).

**(B)** Correlation between the median percent spliced-in (PSI) reported across 56 tissues in the ASCOT database and the adjusted MFE across exons of different strengths, among skippable exons flanked by non-inverted Alu pairs. SS: strong 3’ acceptor, strong 5’ donor; WS: weak 3’ acceptor, strong 5’ donor; SW: strong 3’ acceptor, weak 5’ donor; WW: weak 3’ acceptor, weak 5’ donor. Bonferroni-adjusted linear regression p-values.

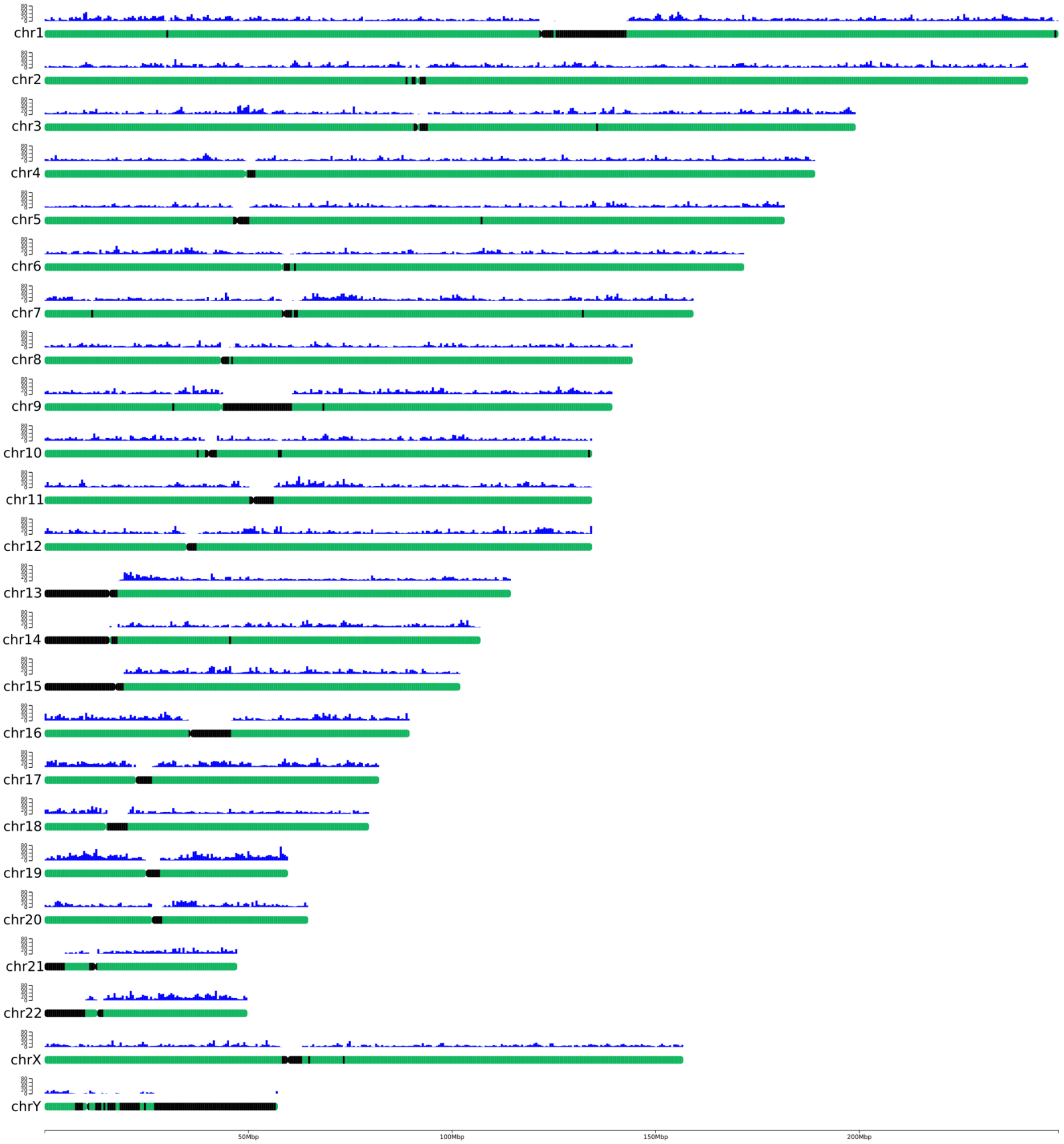


**Figure S3.** Genome-wide distribution of hominoid-specific Alus across all 24 human chromosomes. Each chromosome is represented by a horizontal bar, with the short arm (p) and long arm (q) indicated. The blue histogram above each chromosome illustrates the density of hominoid-specific Alu insertions. Regions that hinder cross-species analysis are colored in black, including pseudo-homologous regions (PHRs) on short arms of acrocentric chromosomes (13, 14, 15, 21, and 22) and pericentromeric regions, as well as the rapidly evolving regions of chromosome Y among primates.

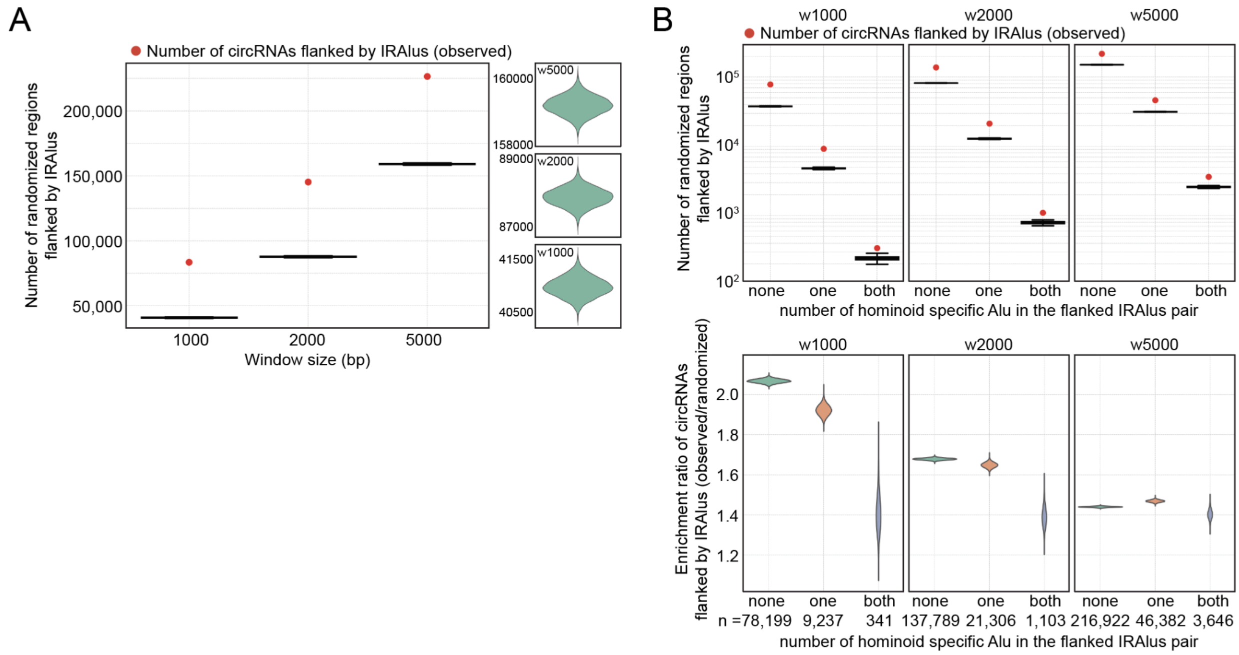


**Figure S4.** Inverted Alu is enriched near circular RNAs.

**(A)** Comparison of the number of circRNAs or randomized regions flanked by IRAlus with the specified window sizes. The observed value is indicated in red. Violin plots showing the detailed distribution for randomized regions are shown together on the right. Region randomization was performed for 10,000 times to ensure robust randomization.

**(B)** Comparison of the number of circRNAs or randomized regions flanked by IRAlus with the specified window sizes and the number of hominoid-specific Alu within the IRAlu pair. The relative enrichment ratios of the circRNAs (observed/randomized) flanked by hominoid-specific IRAlus are shown together in the below. Region randomization was performed for 10,000 times to ensure robust randomization.

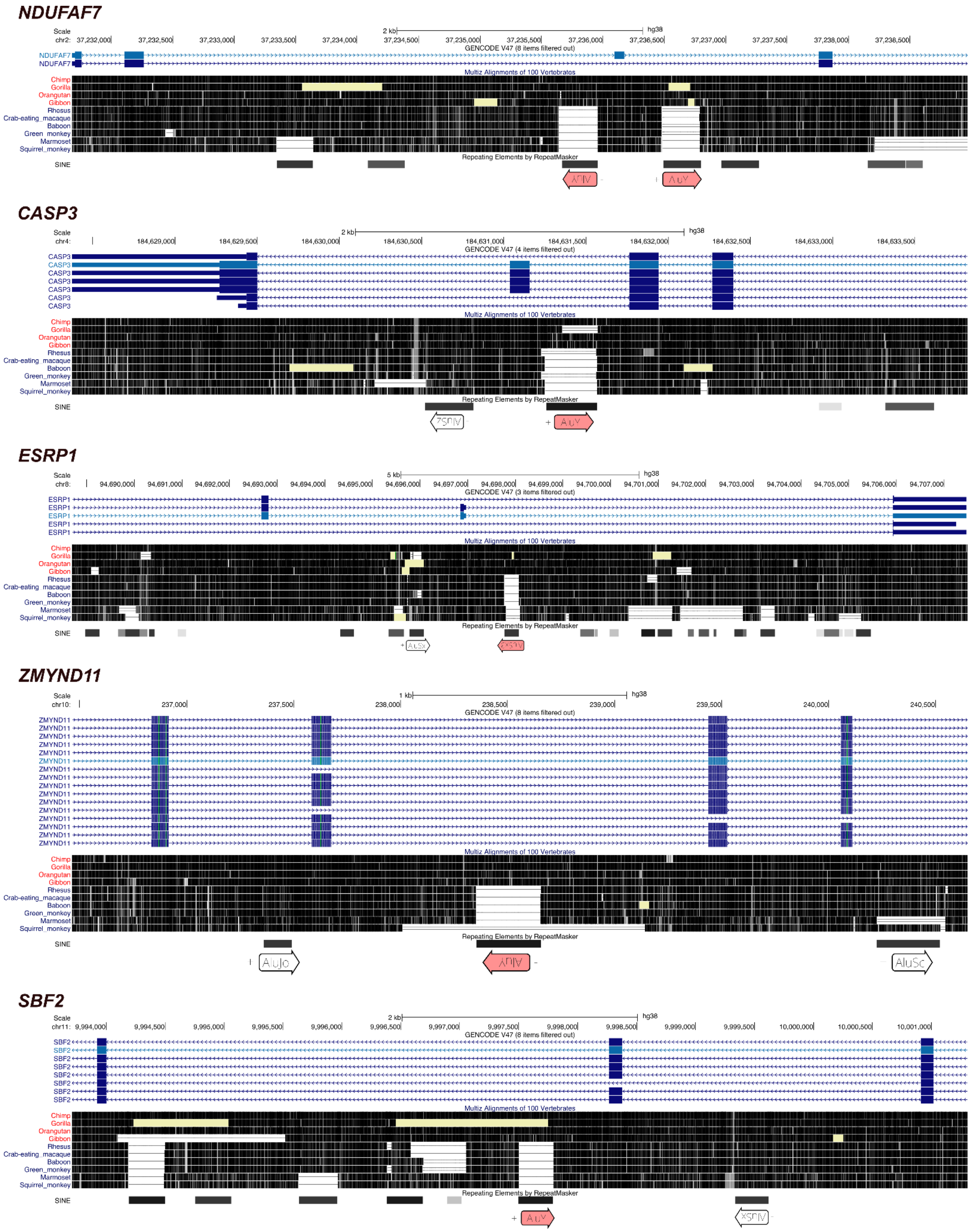


**Figure S5.** UCSC Genome Browser displays of validation candidates.

### *CEP290*

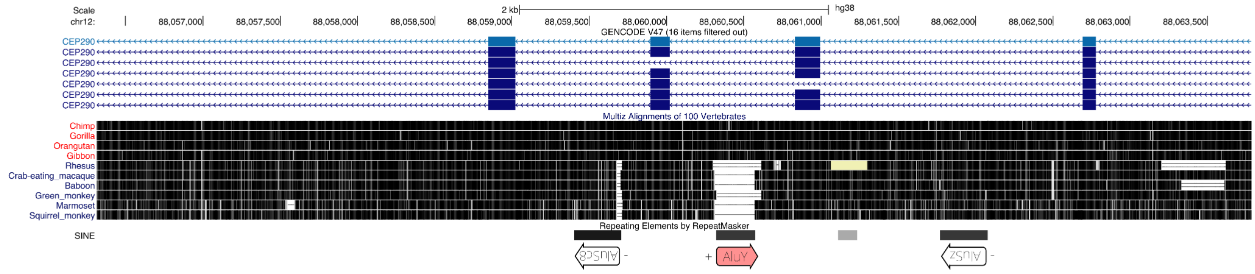


### *DNAH9*

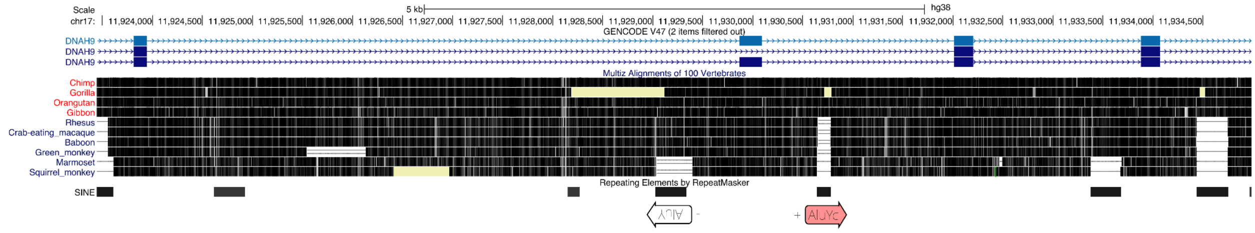


### *PKNOX1*

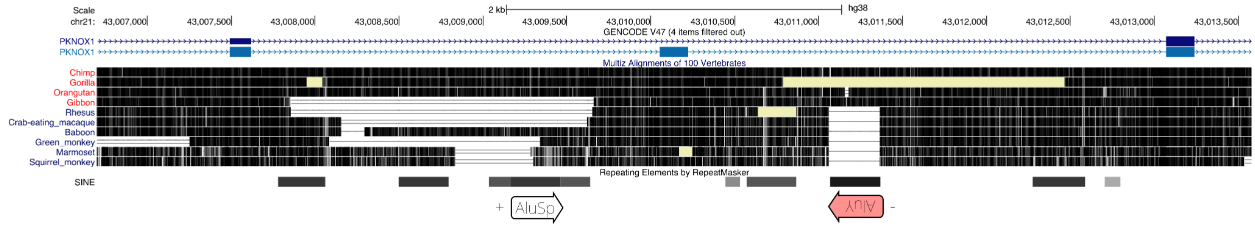


### *CRYBB3*

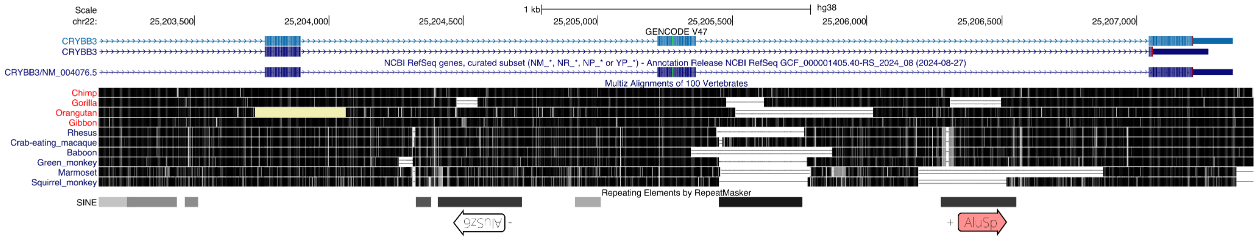


**Figure S5 (Continued).** UCSC Genome Browser displays of validation candidates.

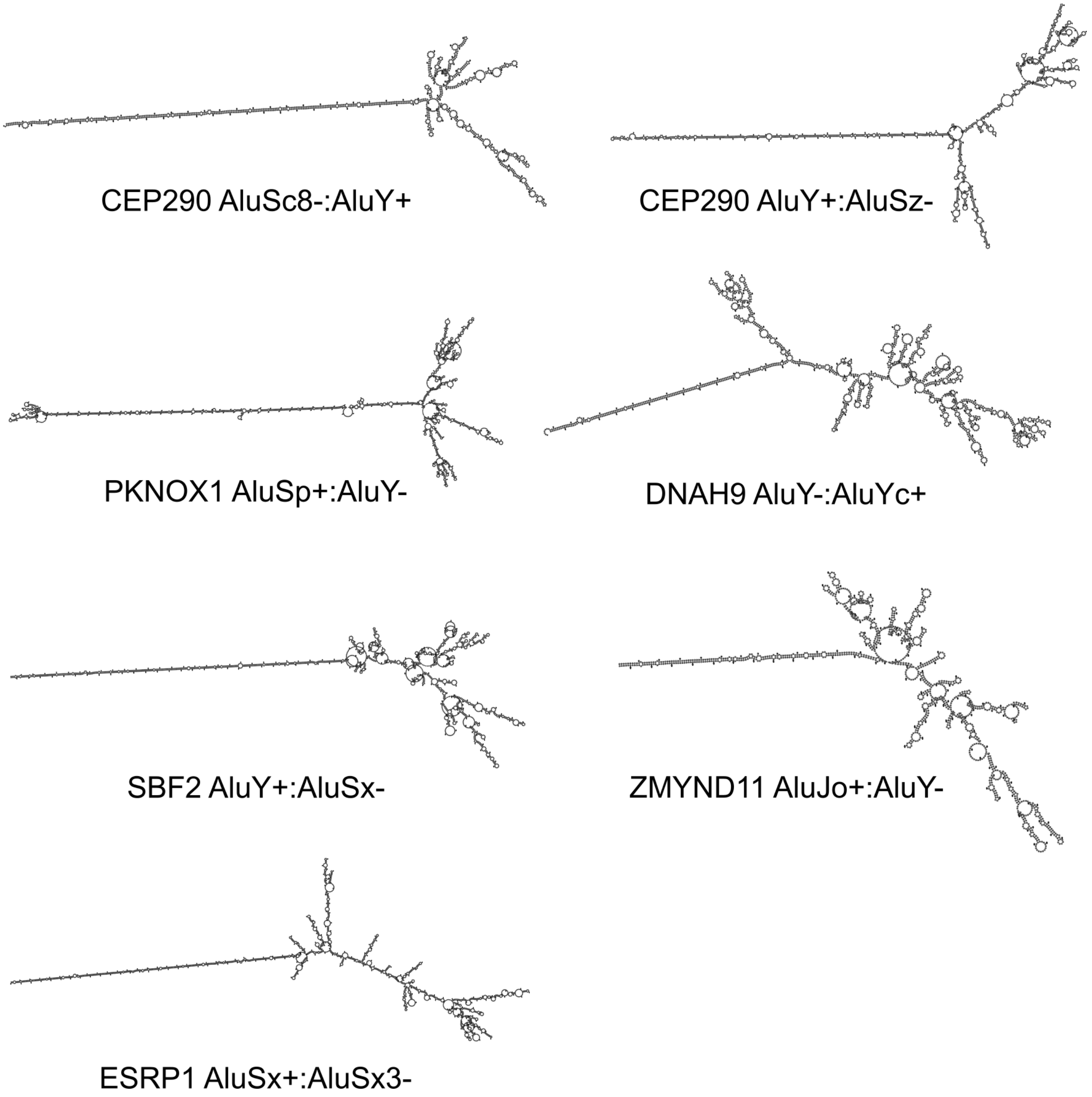


**Figure S6.** RNA secondary structure predictions of candidates for experimental validation.
